# Microglia Depletion Reduces Human Neuronal APOE4-Driven Pathologies in a Chimeric Alzheimer’s Disease Model

**DOI:** 10.1101/2023.11.10.566510

**Authors:** Antara Rao, Nuo Chen, Min Joo Kim, Jessica Blumenfeld, Oscar Yip, Yanxia Hao, Zherui Liang, Maxine R. Nelson, Nicole Koutsodendris, Brian Grone, Leo Ding, Seo Yeon Yoon, Patrick Arriola, Yadong Huang

## Abstract

Despite strong evidence supporting the involvement of both apolipoprotein E4 (APOE4) and microglia in Alzheimer’s Disease (AD) pathogenesis, the effects of microglia on neuronal APOE4-driven AD pathogenesis remain elusive. Here, we examined such effects utilizing microglial depletion in a chimeric model with human neurons in mouse hippocampus. Specifically, we transplanted homozygous APOE4, isogenic APOE3, and APOE-knockout (APOE-KO) induced pluripotent stem cell (iPSC)-derived human neurons into the hippocampus of human APOE3 or APOE4 knock-in mice, and depleted microglia in half the chimeric mice. We found that both neuronal APOE and microglial presence were important for the formation of Aβ and tau pathologies in an APOE isoform-dependent manner (APOE4 > APOE3). Single-cell RNA-sequencing analysis identified two pro-inflammatory microglial subtypes with high MHC-II gene expression that are enriched in chimeric mice with human APOE4 neuron transplants. These findings highlight the concerted roles of neuronal APOE, especially APOE4, and microglia in AD pathogenesis.

**HIGHLIGHTS:**

- Transplanted human APOE4 neurons generate Aβ and p-tau aggregates in APOE4-KI mouse hippocampus.
- Human neuronal APOE4 promotes the formation of dense-core Aβ plaques and p-tau aggregates.
- Microglia is required for human neuronal APOE4-driven formation of p-tau aggregates.
- scRNA-seq reveals enrichment of MHC-II microglia in mice with human APOE4 neuron transplants.

## INTRODUCTION

Apolipoprotein E4 (*APOE4*) is the strongest genetic risk factor for Alzheimer’s Disease (AD). Of the three main APOE isoforms—APOE2, APOE3, and APOE4—APOE4 increases AD risk, reduces age of disease onset, and promotes classical AD pathologies, including deposition of amyloid-beta (Aβ) peptides and hyperphosphorylated tau (p-tau) protein^1–7^. Although APOE4 is strongly linked to AD^1,2,8,9^, its roles in AD pathogenesis are complicated and merit continued exploration. While astrocytes are the main producers of APOE in the central nervous system, APOE can be produced by neurons and microglia in response to stress, injury, or aging^10–16^. Studies have shown that APOE4 has different AD-relevant effects depending on the cell type in which it is produced^4,6,17^.

Several studies have documented the detrimental effects of APOE4 in AD, using either mouse models or human induced pluripotent stem cell (hiPSC)-derived cellular models. Mouse models have traditionally provided a convenient system for studying the multifarious AD-related effects of APOE4 in a diverse, mature *in vivo* environment. In human APOE3 knock-in (E3KI) or APOE4 knock-in (E4KI) mice expressing mutant human amyloid precursor protein (APP) or tau, several experiments have demonstrated that APOE4 exacerbates the formation of Aβ plaques and tau tangles and promotes neurodegeneration^18–20^. In particular, neuronal APOE4 has proved to significantly affect AD pathology^16,21^. However, these mouse model studies have limitations in therapeutic translatability, as they often rely on early-onset AD or other tauopathy-related mutations to produce pathologies and lack key human-specific cellular hallmarks of late-onset AD^6,22^. To study human-specific aspects of late-onset AD, many scientists have turned to *in vitro* hiPSC-derived cell models. Several labs using hiPSC-derived neurons have identified cellular features of APOE4-driven AD pathologies, including increased levels of secreted Aβ species and intracellular phosphorylated tau species^12,23,24^. However, these *in vitro* hiPSC-derived models are restricted in their ability to model mature cell behavior, produce Aβ or tau aggregates, and recapitulate multi-cellular interactions as seen *in vivo*.

To address these limitations, our lab developed an *in vivo* chimeric late-onset AD mouse model comprised of hiPSC-derived neurons with different APOE genotypes transplanted into APOE knock-in (EKI) mouse hippocampi^25^. This hybrid model facilitates human neuronal maturation and AD pathology far beyond what is capable *in vitro*, even producing Aβ aggregates reminiscent of pathology found in late-onset AD^25^. In addition, this model allows for investigation of how human neurons interact with other cell types *in vivo* in APOE4-related AD pathogenesis. We were particularly interested in microglia, which have been consistently implicated in many aspects of AD pathogenesis^26–28^. Microglia play key roles in maintaining a healthy brain environment, including surveillance of and activated response to cellular debris as well as phagocytosis of pathological protein aggregates, like Aβ and tau assemblies^29,30^. Recent studies have found that microglia in AD may interact with neurons to contribute to AD pathogenesis, either by reducing clearance of Aβ and tau^7,23,31,32^ or by active seeding/spreading of Aβ and tau aggregates^28,33–35^. The recent development of effective microglial depletion tools has allowed us to begin distinguishing between these two possibilities in the context of different APOE isoforms. In particular, PLX3397 (PLX) is a potent inhibitor of vital microglial protein colony-stimulating factor 1 receptor (Csf1r), allowing for depletion of microglia without significantly affecting other cell types in the brain^36,37^. We used PLX depletion of microglia in our chimeric model mice to investigate the roles of microglia in modifying neuronal APOE4-driven AD pathologies.

## RESULTS

### Transplanted hiPSC-derived neurons survive in APOE-KI mouse hippocampus

Our lab previously generated and characterized three different hiPSC lines: 1) homozygous human APOE4 (hE4) hiPSCs from an AD patient, 2) isogenic human APOE3 (hE3) hiPSCs generated from the E4 hiPSC line by gene editing, and 3) human APOE-knockout (hEKO) hiPSCs from a patient homozygous for an ablative *APOE* frameshift mutation (c.291del, p.E97fs)^12,38^. To generate a chimeric AD model, we differentiated these three hiPSC lines into mixed neuronal cultures (including both excitatory and inhibitory neurons) using an established neuronal differentiation protocol^25^, and transplanted them into the hippocampi of 4-month-old E4KI and E3KI mice. hE4 and hE3 neurons were transplanted into E4KI and E3KI mice of matching APOE genotype (hE4 neurons into E4KI mice and hE3 neurons into E3KI mice), while hEKO neurons were transplanted into E4KI mice to examine the interactive effects of human neurons lacking APOE with APOE4-expressing microglia. To improve cell transplant survival, mouse host immune response was blocked with an immunosuppressant cocktail administered every other day for a week after transplantation (doses on days 0, 2, 4, and 6)^39^. The chimeric mice were then aged for 8 months, during which time they were either fed an AIN-76A control chow for the entire duration or fed a control chow for the first 4 months and then fed an AIN-76A-PLX chow containing 400 mg/kg of PLX for the remaining 4 months (Figure 1A). Conditions were labeled according to the *APOE* genotype of the transplanted human cells (hE3-, hE4-, hEKO-), the *APOE* genotype of the host mice (E3KI, E4KI), and the type of chow the chimeric mice received (control chow, PLX chow) (Figure 1A). In total, six groups of mice were used in this study: hE4-E4KI, hE4-E4KI-PLX, hE3-E3KI, hE3-E3KI-PLX, hEKO-E4KI, hEKO-E4KI-PLX (Figure 1A).

**Figure 1.**
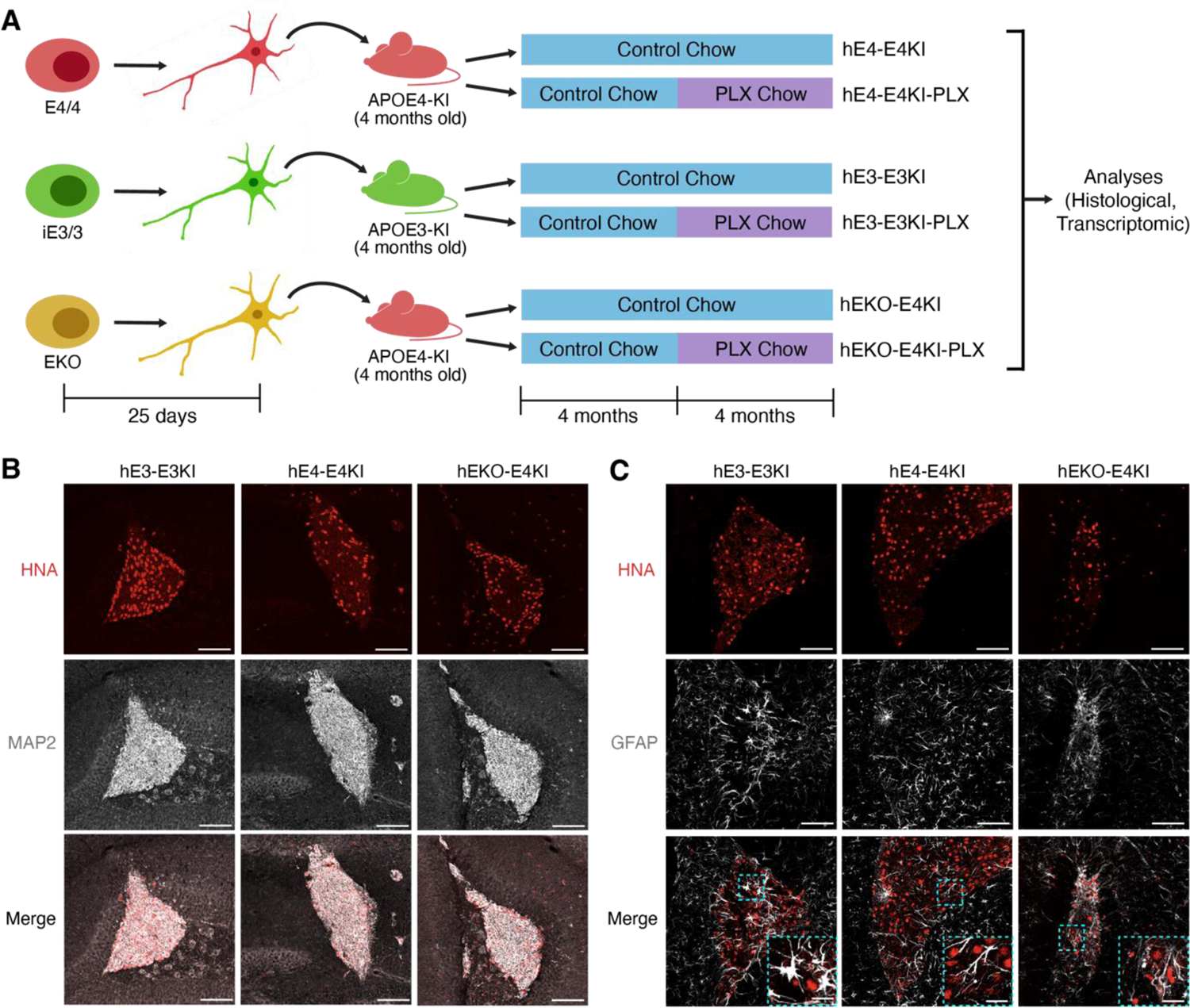
Transplanted human neuronal progenitors survive and develop into neurons in chimeric mouse hippocampus. **(A)** Experimental design: iPSCs with different APOE genotypes were differentiated into neuronal progenitors and transplanted into human APOE-KI mice. The chimeric mice were aged for 8 months, with half the mice receiving PLX3397 (PLX) for the latter 4 months. Conditions were labeled accordingly, and all mice were used for histological or transcriptomic analysis. **(B)** Representative immunostaining images of the human cell transplants in the hippocampus of the 12-month-old chimeric mice (8 months post transplantation). Top row: Human Nuclear Antigen (HNA, red) to mark the human cell transplants; second row: human-preferential MAP2 (gray) to mark neurons; third row: composite of HNA and MAP2 images. Scale bar, 100 μm. **(C)** Representative immunostaining images of the human cell transplants in the hippocampus of the 12-month-old chimeric mice. Stained with HNA (red) for human cell transplants and GFAP (gray) for astrocytes. Top row: Human Nuclear Antigen (HNA, red) to mark the human cell transplants; second row: GFAP (gray) to mark astrocytes; third row: composite of HNA and GFAP images, with magnified image insets showing that HNA^+^ cells and GFAP^+^ cells do not overlap. Scale bar, 100 μm. Scale bar for magnified image insets, 25 μm.

At 8 months post-transplantation, the chimeric mice were transcardially perfused, with one brain hemisphere fixed for immunohistochemical analyses and the other hemisphere harvested for single cell RNA-sequencing (scRNA-seq) of isolated hippocampal microglia. Transplanted cells stained positive for human nuclear antigen (HNA) (Figures 1B and 1C) and for mature neuronal marker MAP2 (Figure 1B). These same HNA^+^ cells did not show positivity for astrocyte marker GFAP (Figure 1C), indicating no astrocytes developed from the transplanted human cells. Together, these data signify successful survival and maturation of the transplanted human neurons in E3KI and E4KI mouse hippocampi. The MAP2 antibody used for these experiments shows preferential recognition of human versus mouse neurons, resulting in the transplanted human neurons appearing substantially brighter than the surrounding mouse neurons (Figure 1B). As a result, for all following stains, we were able to use MAP2 staining to identify the transplanted human neurons.

### PLX treatment effectively depletes microglia, but not astrocytes, in the hippocampus of chimeric mice

We then measured the effects of PLX treatment on microglia counts in the hippocampus. Upon staining the chimeric mouse brain sections with microglial marker Iba1, we found significant depletion of microglia in all three groups treated with PLX (Figure 2A and 2B). Unexpectedly, microglial depletion was more efficacious in hE4-E4KI-PLX mouse hippocampus (93%) than in hEKO-E4KI-PLX (84%) and hE3-E3KI-PLX (67%) mouse hippocampus. While this attenuated microglial depletion effect in hE3-E3KI-PLX mice may be enhanced by the increased baseline number of microglia in hE3-E3KI mice compared to hE4-E4KI mice (Figure 2B), this difference merited further investigation.

**Figure 2.**
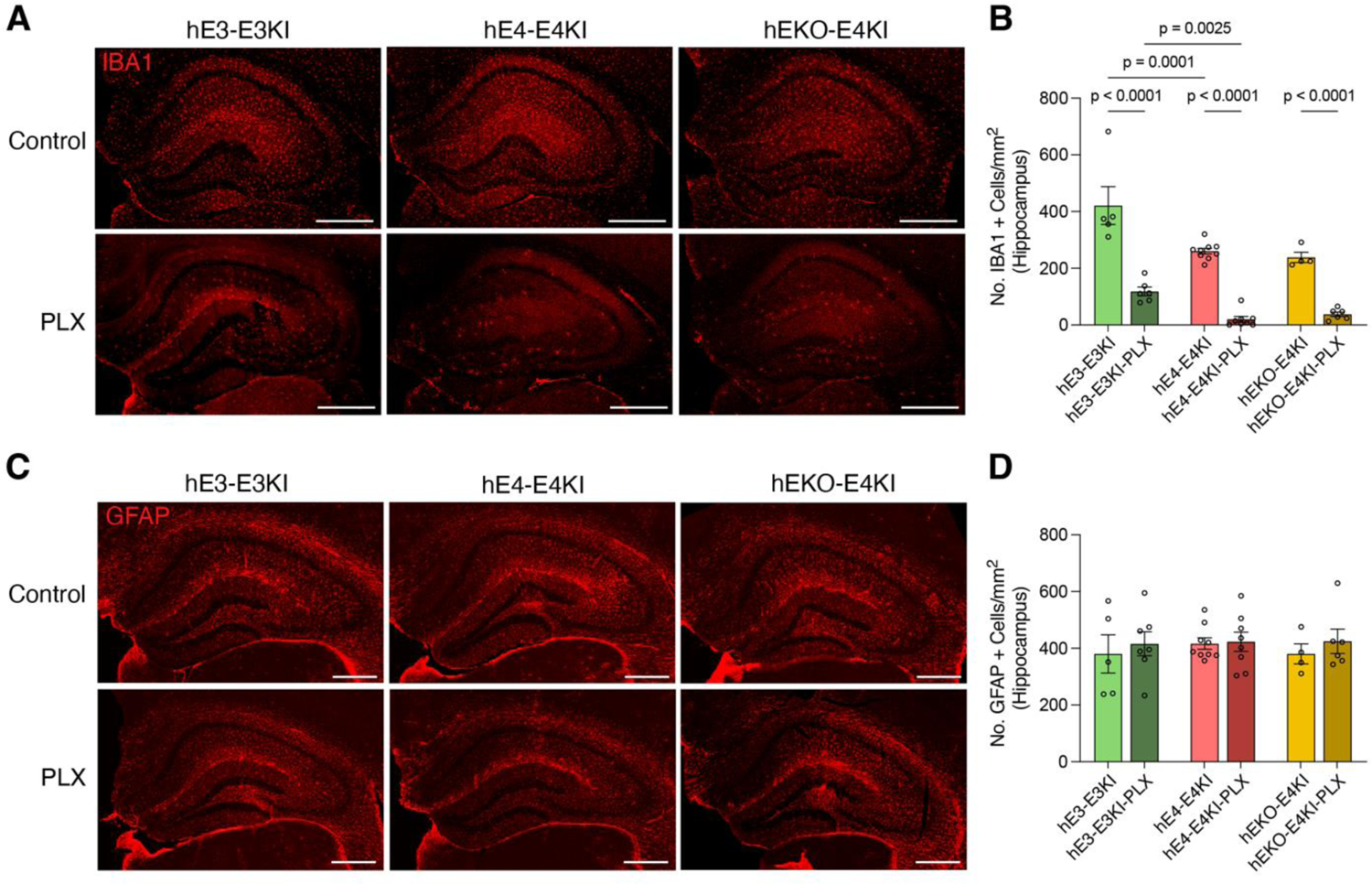
PLX depletes microglia, but does not affect astrocytes, in chimeric mouse hippocampus. **(A)** Representative immunostaining images of microglia (Iba1, red) in the hippocampus of chimeric mice of each condition. Scale bar, 500 μm. **(B)** Quantification of number/mm^2^ of Iba1^+^ microglia in the hippocampus. Each dot represents one mouse per condition (hE3-E3KI, n=5; hE3-E3KI-PLX, n=7; hE4-E4KI, n=9; hE4-E4KI-PLX, n=8; hEKO-E4KI, n=4; hEKO-E4KI-PLX, n=6). **(C)** Representative immunostaining images of astrocytes (GFAP, red) in the hippocampus of chimeric mice of each condition. Scale bar, 500 μm. **(D)** Quantification of number/mm^2^ of GFAP^+^ astrocytes in the hippocampus. Each dot represents one mouse per condition (hE3-E3KI, n=5; hE3-E3KI-PLX, n=7; hE4-E4KI, n=9; hE4-E4KI-PLX, n=8; hEKO-E4KI, n=4; hEKO-E4KI-PLX, n=6). All data are expressed as mean ± S.E.M. Differences between groups were determined by two-way ANOVA with Benjamini’s post hoc test for multiple comparisons.

To this end, we sought to measure baseline levels of Csf1r expression in microglia of E3KI and E4KI mouse hippocampus. To match the age at which PLX was first administered, we took 8-month-old untransplanted E3KI and E4KI mice, isolated hippocampal CD11b^+^/CD45^int^ microglia using fluorescence-activated cell sorting (FACS), and measured Csf1r levels on a BD LSR Fortessa™ X-20 cell analyzer (Figure S1A). When comparing each of the six mice per condition (Figure S1B), we found that E4KI microglia showed significantly increased (38%) median fluorescence intensity (MFI) of Csf1r than E3KI microglia (Figures S1C and S1D). This suggests that higher baseline expression of Csf1r, which may be an APOE4-driven effect in microglia, renders microglia more susceptible to the effects of Csf1r antagonist PLX.

We also tested whether PLX depleted astrocytes, the other main glial cell type in the hippocampus. Staining with astrocyte marker GFAP revealed no significant differences in hippocampal astrocyte number across all chimeric mouse conditions (Figures 2C and 2D). This result is in line with other studies that have reported no changes in overall astrocyte viability upon PLX dosage both *in vitro* and *in vivo*^37,40,41^.

### Microglial depletion decreases Aβ aggregates in the presence of neuronal APOE4

We previously reported that transplanted human neurons produced Aβ aggregates and Thioflavin-S^+^ small plaques in EKI mice, with APOE4 exacerbating this phenotype^25^. The ability of transplanted human neurons to generate Aβ aggregates, which do not form in human *in vitro* neuronal models^12,42^ nor in mouse *in vivo* models without familial AD mutations^43,44^, provided us a unique opportunity to study the impact of microglia with different APOE genotypes on human neuronal APOE-driven Aβ pathologies, which better mimic late-onset AD. When we stained the chimeric mouse hippocampal sections with human Aβ-specific monoclonal antibody 3D6, we found 3D6^+^ Aβ aggregates formed within or immediately surrounding the MAP2^+^ transplanted human neuronal cells in all conditions (Figure 3A). Similarly to our earlier study with a chimeric mouse model^25^, a significant majority of these Aβ aggregates formed within 100 μm of the human neuron transplants (Figure S2A, S2C, and S2E) and very few in hippocampal regions outside that boundary (Figure S2F). These replicated results reiterate the conclusion that the human neuronal transplants are the source of these Aβ aggregates.

**Figure 3.**
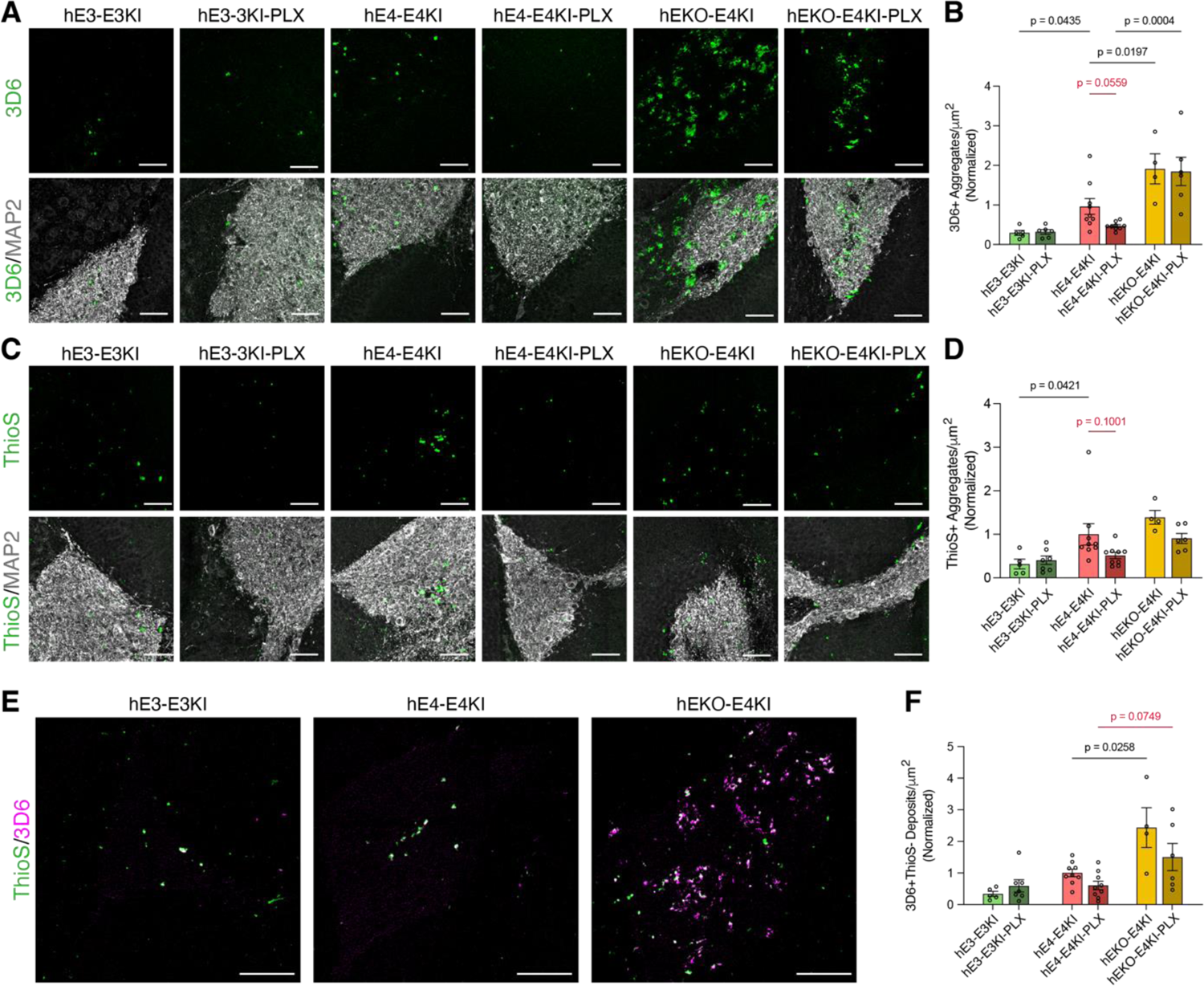
Human neuronal APOE isoform and microglial depletion affect Aβ pathology in chimeric mouse hippocampus. **(A)** Representative immunohistochemical images of Aβ aggregates within and immediately surrounding human neuronal transplants. Top row: 3D6^+^ Aβ aggregates (green); second row: composite of 3D6^+^ Aβ aggregates (green) and MAP2^+^ (gray) human neuron transplants. Scale bar, 50 μm. **(B)** Quantification of number of 3D6^+^ Aβ aggregates/μm^2^ within a 100 μm perimeter per transplant area (hE3-E3KI, n=5; hE3-E3KI-PLX, n=6; hE4-E4KI, n=9; hE4-E4-KI-PLX, n=9; hEKO-E4KI, n=4; hEKO-E4KI-PLX, n=6). **(C)** Representative immunohistochemical images of Thioflavin-S^+^ dense-core Aβ aggregates within and immediately surrounding human neuronal transplants. Top row: Thioflavin-S^+^ dense-core Aβ aggregates (green); second row: composite of Thioflavin-S^+^ dense-core Aβ aggregates (green) and MAP2^+^ (gray) human neuron transplants. Scale bar, 50 μm. **(D)** Quantification of number of Thioflavin-S^+^ dense-core Aβ aggregates/μm^2^ within a 100 μm perimeter per transplant area (hE3-E3KI, n=5; hE3-E3KI-PLX, n=7; hE4-E4KI, n=9; hE4-E4KI-PLX, n=9; hEKO-E4KI, n=4; hEKO-E4KI-PLX, n=6). **(E)** Representative immunohistochemical images of 3D6^+^ Aβ aggregates (magenta) and Thioflavin-S^+^ dense-core Aβ aggregates (green) within human neuron transplants. Diffuse Aβ aggregates were defined as 3D6^+^/Thioflavin-S^-^ aggregates. Scale bar, 100 μm. **(F)** Quantification of number of diffuse (3D6^+^/Thioflavin-S^-^) Aβ deposits/μm^2^ within a 100 μm perimeter per transplant area (hE3-E3KI, n=5; hE3-E3KI-PLX, n=7; hE4-E4KI, n=9; hE4-E4KI-PLX, n=9; hEKO-E4KI, n=4; hEKO-E4KI-PLX, n=6). For all quantifications, values are normalized to the hE4-E4KI condition, and each dot represents one mouse per condition. All data are expressed as mean ± S.E.M. Differences between groups were determined by two-way ANOVA with Benjamini’s post hoc test for multiple comparisons. Comparisons of P<0.05 were considered significant and are displayed in black. Comparisons of 0.05<P<0.11 are displayed in red.

For all following quantifications, we compared: 1) hE4-E4KI mice versus hE3-E3KI mice and hE4-E4KI-PLX mice versus hE3-E3KI-PLX mice to measure the effects of APOE4 versus APOE3, 2) hEKO-E4KI mice versus hE4-E4KI mice and hEKO-E4KI-PLX mice versus hE4-E4KI-PLX mice to specifically measure the effects of neuronal APOE-KO versus APOE4, and 3) mice fed control chow versus PLX chow within each APOE genotype group to measure the effects of microglial depletion. All conditions were normalized to results from hE4-E4KI mice, since they served as the common factor in comparisons with both hE3-E3KI and hEKO-E4KI mice.

When we quantified the number of Aβ aggregates per square micrometer within 100 μm of the transplants, we found that APOE isoform had a significant effect on Aβ aggregate counts. Of the chimeric mice fed with control chow, the hE3-E3KI mice displayed significantly fewer Aβ aggregates than hE4-E4KI mice (Figure 3B), confirming previous findings that APOE4 exacerbates amyloid pathology^12,19,25,45^. Upon microglial depletion, the average number of Aβ aggregates in the hE4-E4KI condition was reduced by half, with a p-value just shy of significance (Figure 3B, red text). These results suggest that the presence of microglia promotes neuronal APOE4-driven Aβ pathology, in line with several studies showing that microglia play a role in seeding amyloid plaques^28,33–35^. In contrast, depleting microglia had no significant effect on number of Aβ aggregates in the hE3-E3KI-PLX condition. Taken together, these data support the conclusion that the presence of both APOE4 neurons and APOE4 microglia promote Aβ aggregate formation.

### Neuronal APOE deficiency increases diffuse Aβ deposits independent of microglia

Furthermore, we initially found that hEKO-E4KI mice had a higher number of 3D6^+^ Aβ aggregate-like deposits than hE4-E4KI mice (Figure 3B). And unlike in hE4-E4KI-PLX mouse hippocampus, microglial depletion had no effect on the number of Aβ deposits in the hEKO-E4KI-PLX mouse hippocampus (Figure 3B). To investigate this effect further, we compared the Aβ in hEKO-E4KI mice, both with and without PLX treatment, to hE4-E4KI mice with neuronal APOE4, and noticed distinct morphological differences in the hEKO-E4KI Aβ aggregate-like deposits, including increased size (Figure S3A) and a diffuse feathered structure (Figure 3A). To test the nature of these deposits, we stained the chimeric mouse hippocampus with dense-core plaque marker Thioflavin-S (Figure 3C). As with Aβ, Thioflavin-S staining showed a significant increase in number of dense-core deposits in hE4-E4KI mice compared to hE3-E3KI (Figure 3D). However, hEKO-E4KI and hE4-E4KI conditions showed no differences in size or number of dense-core deposits (Figure 3D, Figure S3B), nor was there a significant reduction in dense-core deposits (although there appeared to be a trend in hE4-E4KI-PLX mice) upon removal of microglia. Importantly, while many Aβ aggregates colocalized with Thioflavin-S in a manner reminiscent of dense-core amyloid plaques in hE3-E3KI and hE4-E4KI mice, the majority of the aggregates stained positive for Aβ were negative for Thioflavin-S in hEKO-E4KI mice (Figure 3E). We quantified these 3D6^+^/Thioflavin-S^-^ Aβ deposits, referred to as “diffuse” Aβ deposits, and saw a significant increase in the number of diffuse Aβ deposits in hEKO-E4KI compared to hE4-E4KI mice (Figure 3F). These results indicate that removal of APOE from human neurons *in vivo* specifically promotes the formation of diffuse Aβ deposits, which are not significantly influenced by microglial depletion. It should be noted that this phenotype occurred in the presence of high levels of astrocyte-secreted APOE4 in hEKO-E4KI mouse hippocampus, indicating that the transplanted human neuron-produced APOE4 is a key driving factor for the formation of dense-core Aβ plaques.

### Microglial depletion reduces p-tau aggregates in the presence of neuronal APOE4

We next examined the chimeric mice for formation of p-tau pathology. Upon staining with p-tau-specific monoclonal antibody AT8, we found that these chimeric mice did indeed produce AT8^+^ p-tau aggregates (Figure 4A), reminiscent of p-tau pathology previously described in a chimeric mouse model of transplanted human neurons with a pathogenic tau mutation^46^. A highly significant majority of these p-tau aggregates, as with the findings for Aβ aggregates, were located within 100 μm of the human neuronal transplants (Figures S2B, S2D, and S2G). In regions in the hippocampus further than 100 μm from the transplant, there were rarely p-tau aggregates found (Figure S2H). Detailed examination of p-tau aggregates seemed to indicate an intracellular localization, with aggregates immediately surrounded by soma-like MAP2^+^ staining (Figure 4B). Taken together, these data strongly suggest that the p-tau aggregates originate from the human neuron transplants.

**Figure 4.**
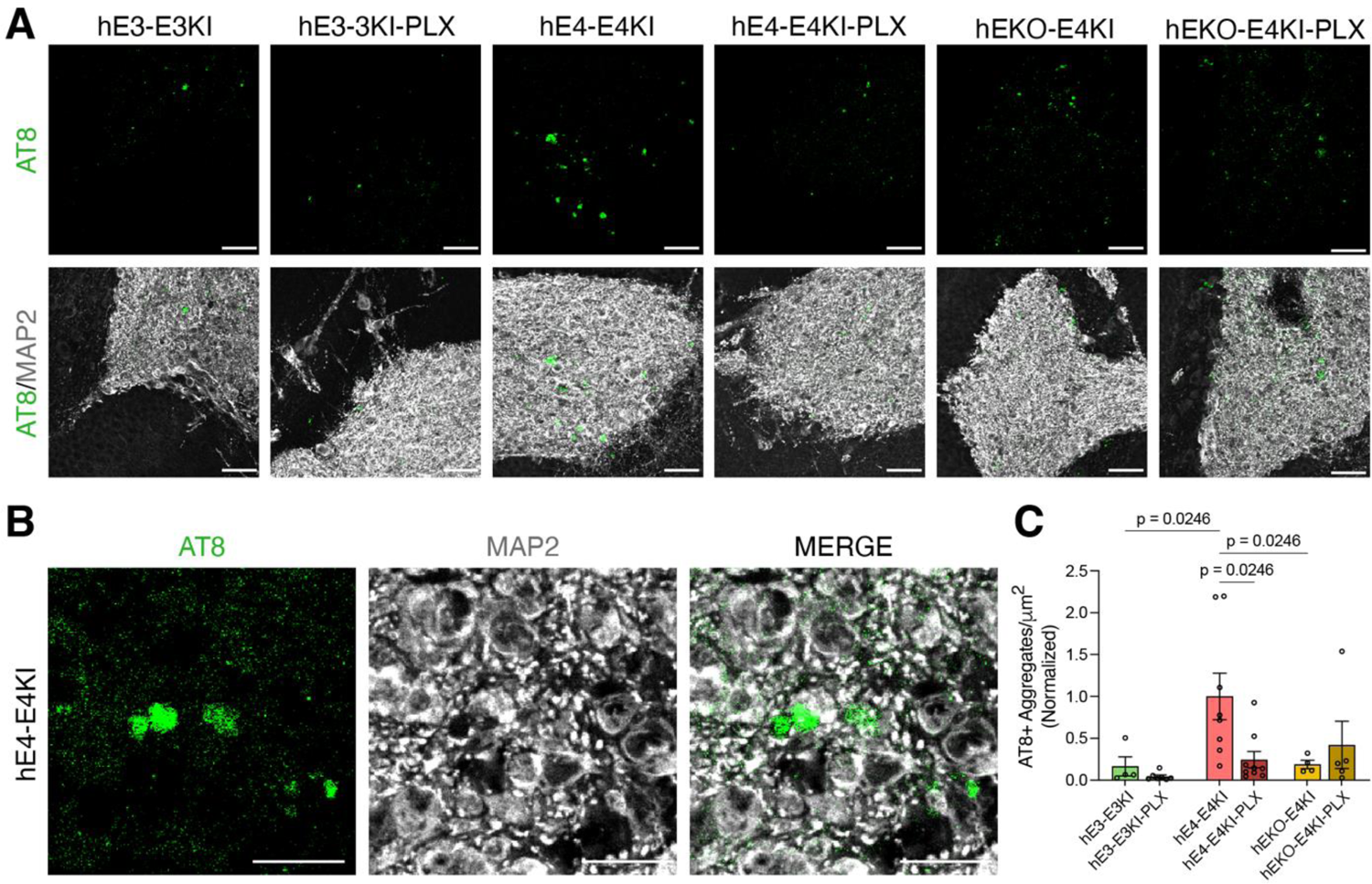
Human neuronal APOE isoform and microglial depletion affect p-tau pathology in chimeric mouse hippocampus. **(A)** Representative immunohistochemical images of p-tau aggregates within and immediately surrounding human neuronal transplants. Top row: AT8^+^ p-tau aggregates (green); second row: composite of AT8^+^ p-tau aggregates (green) and MAP2^+^ (gray) human neuron transplants. Scale bar, 50 μm. **(B)** Many AT8^+^ p-tau aggregates are present within transplanted MAP2^+^ human neurons. Each image represents the same field of view, stained for p-tau (AT8, green, first column), human neuronal transplants (MAP2, gray, second column), and a composite of AT8 and MAP2 (third column). Scale bar, 20 μm. **(C)** Quantification of number of AT8^+^ p-tau aggregates/μm^2^ within a 100 μm perimeter per transplant area. Values are normalized to the hE4-E4KI condition, and each dot represents one mouse per condition (hE3-E3KI, n=4; hE3-E3KI-PLX, n=7; hE4-E4KI, n=8; hE4-E4KI-PLX, n=9; hEKO-E4KI, n=4; hEKO-E4KI-PLX, n=5). All data are expressed as mean ± S.E.M. Differences between groups were determined by two-way ANOVA with Benjamini’s post hoc test for multiple comparisons.

When we quantified the number of p-tau aggregates per square micrometer within 100 μm of the transplants, we found that the average number of aggregates in the hE4-E4KI mouse hippocampus was significantly higher than in both the hE3-E3KI and hEKO-E4KI mouse hippocampus. The reduced p-tau pathology in human neuronal APOE3 and APOE-KO conditions matches a recently published study showing that, compared to APOE4, APOE3 and neuronal APOE4-KO can dramatically reduce tau pathology in PS19/EKI mice^21^. Upon microglial depletion, hE4-E4KI-PLX mouse hippocampus had significantly reduced levels of p-tau aggregates compared to control hE4-E4KI mouse hippocampus (Figure 4C), although aggregate size was unaffected across all conditions (Figure S3C). Interestingly, microglial depletion did not significantly affect p-tau aggregate numbers in hE3-E3KI-PLX or hEKO-E4KI-PLX mouse hippocampus. Taken together, these results further support the conclusion that E4KI microglia promote formation of AD pathology, this time with p-tau aggregates in addition to amyloid, in the presence of human neuronal APOE4.

### Microglial depletion increases APOE levels within human neuron transplants

We then examined whether and how microglial depletion affects APOE levels in human neuron transplants with different APOE genotypes. When we analyzed APOE protein by immunohistochemistry within the human neuron transplants of the control chimeric mice, we found relatively low APOE expression, most of which colocalized with GFAP^+^ astrocytes (Figure 5A). Once microglia were depleted, the APOE staining within the transplants increased significantly in hE3-E3KI-PLX and hE4-E4KI-PLX mice (Figure 5B), suggesting that microglia may play a role in clearing transplanted human neuron-produced APOE. Interestingly, hEKO-E4KI-PLX mice had around half of the APOE staining within the transplants compared to hE4-E4KI-PLX mice, suggesting that at least ∼50% of the transplant-localized APOE4 in hE4-E4KI mice is from the transplanted human neurons. In accordance with this finding, we found that APOE staining in the transplants of hE3-E3KI-PLX and hE4-E4KI-PLX mouse hippocampus was colocalized with both GFAP^+^ astrocytes and MAP2^+^ human neurons (Figure 5A), and APOE staining significantly correlated with human neuron transplant area in hE4-E4KI-PLX mice and moderately in hE3-E3KI-PLX mice (Figure 5F-G). Interestingly, microglia numbers were negatively correlated with intra-transplant APOE staining in hE4-E4KI-PLX mouse hippocampus (Figure S4E), supporting the conclusion that microglia clear the transplanted human neuron-produced APOE. Together, these data indicate that microglia may operate to clear APOE generated from the transplanted human neurons. Neurons with high levels of APOE have previously been shown in PS19/E4KI mice upon microglial depletion^37^, supporting our conclusion.

**Figure 5.**
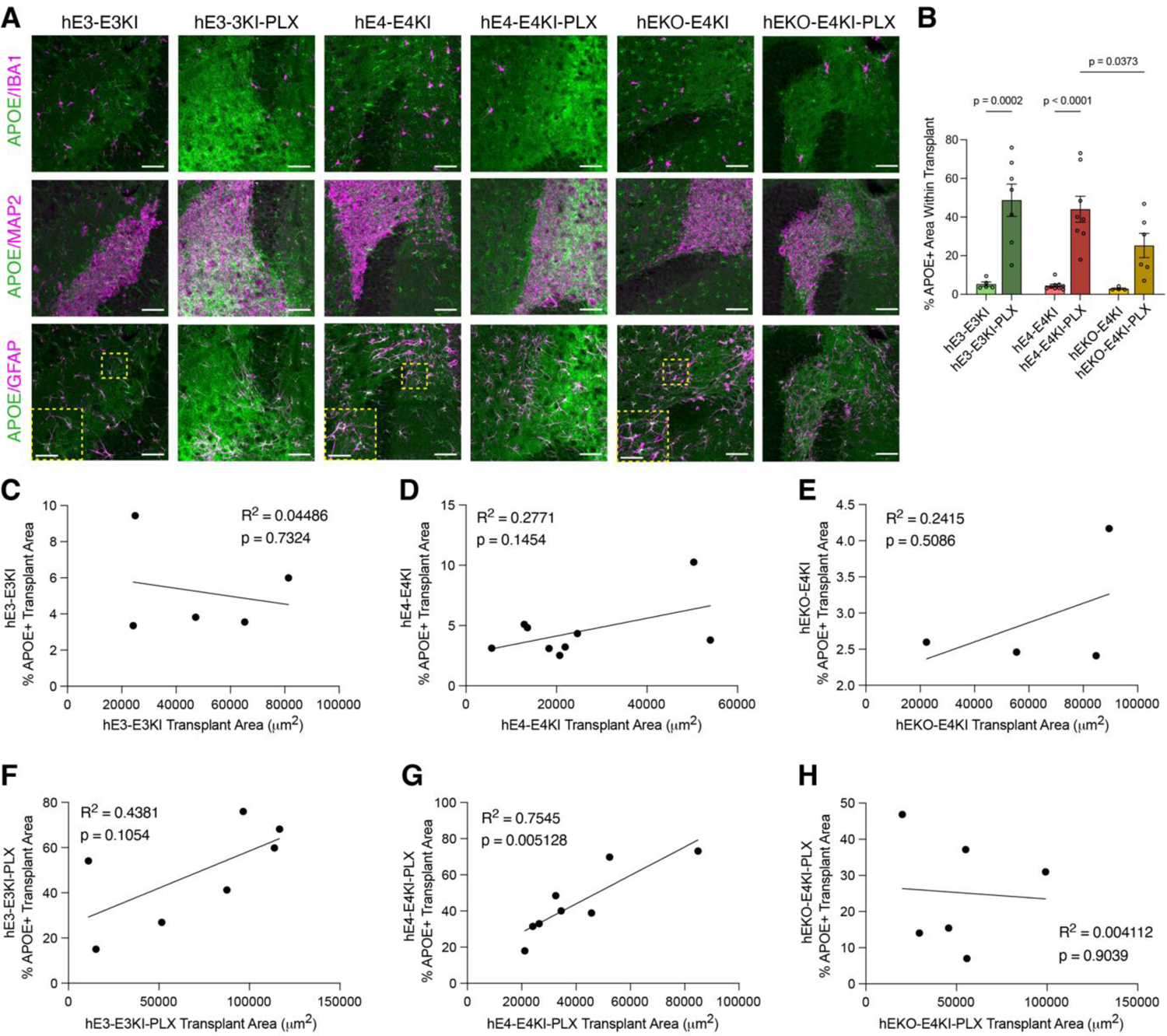
Microglial depletion increases APOE levels within human neuronal transplants. **(A)** Representative immunohistochemical images of APOE within human neuronal transplants. Each column represents the same field of view, with a composite of APOE (green) with Iba1^+^ microglia (magenta, top row), MAP2^+^ human neuron transplants (magenta, second row), and GFAP^+^ astrocytes (magenta, third row, with magnified image insets showing APOE and GFAP overlap). Scale bar, 50 μm. Scale bar for magnified insets, 25 μm. **(B)** Quantification of average APOE fluorescence intensity within the human neuron transplants. Values are normalized to the hE4-E4KI condition, and each dot represents one mouse per condition (hE3-E3KI, n=5; hE3-E3KI-PLX, n=7; hE4-E4KI, n=9; hE4-E4KI-PLX, n=8; hEKO-E4KI, n=4; hEKO-E4KI-PLX, n=6). All data are expressed as mean ± S.E.M. Differences between groups were determined by two-way ANOVA with Benjamini’s post hoc test for multiple comparisons. **(C)** Correlations for each condition between % ApoE^+^ transplant area and size of transplants. Each dot represents one mouse per condition (hE3-E3KI, n=5; hE3-E3KI-PLX, n=7; hE4-E4KI, n=9; hE4-E4KI-PLX, n=8; hEKO-E4KI, n=4; hEKO-E4KI-PLX, n=6). Pearson’s correlation analyses (two-sided).

### Transcriptional characterization of hippocampal microglia in chimeric mice

To investigate microglial reactivity to human neuron transplants in each condition, we performed scRNA-seq on microglia from the hippocampi of chimeric mice. We dissected and dissociated hippocampi from mice of chimeric conditions (hE3-E3KI, hE4-E4KI, hEKO-E4KI) and untransplanted control genotypes (E3KI and E4KI). Microglia were then purified via fluorescence activated cell sorting (FACS) for scRNA-seq (Figure 6A). Flow cytometry gating involved removing debris, non-single cells, and dead cells, before sorting specifically for CD11b^+^/CD45^int^ microglia (Figure S5). After scRNA-seq, we clustered the sequenced cells, filtered for quality control (Figure S6A–S6E), and assessed the purity of the sorted microglia. Cells were clustered by the Louvain algorithm^47^ and visualized by Uniform Manifold Approximation and Projection (UMAP), revealing 16 distinct cell clusters (Figure 6B). Nearly all the sequenced cells expressed microglia-specific markers *Cx3cr1* and *Csf1r* (Figures 6C and 6D), indicating that almost all of the sequenced cells were microglia. Interestingly, expression of *Cd74*, a pro-inflammatory MHC-II marker for reactive microglia^48,49^, highlighted select microglial clusters, including clusters 4 and 12, that warranted further investigation (Figure 6E and see below). When tested for markers of non-microglial cell types, the sequenced cells had very sparse or no expression of *Syn1* (neurons), *Gfap* (astrocytes), *Slc17a7* (excitatory neurons), *Gad1/Gad2* (inhibitory neurons), and *Mbp* (oligodendrocytes) (Figure 6F-K), further supporting the high purity of the sorted microglia.

**Figure 6.**
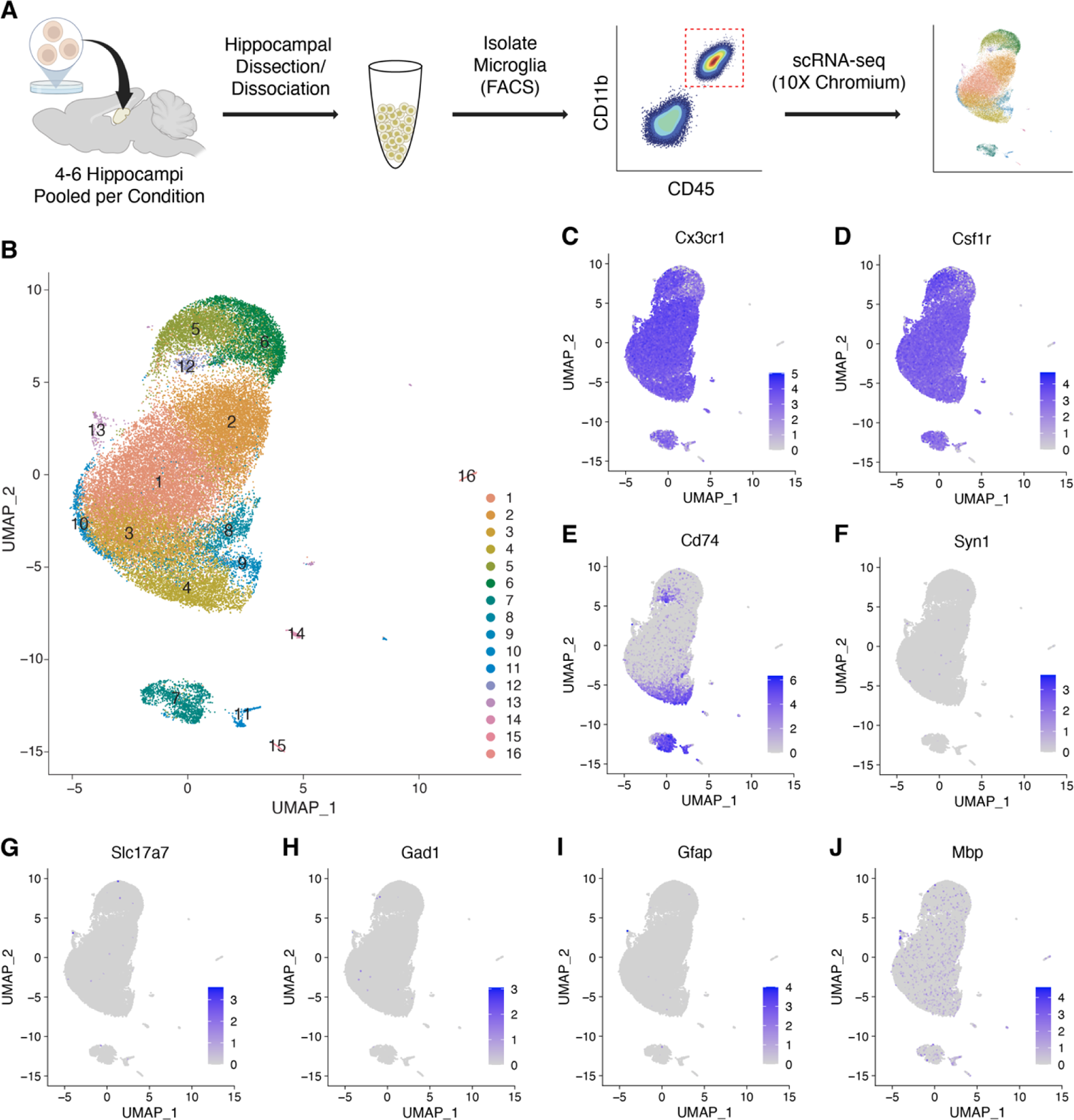
Transcriptional characterization of hippocampal microglia in chimeric mouse hippocampus. **(A)** Workflow summarizing isolation of CD11b^+^/CD45^int^ microglia from the pooled hippocampi of chimeric mice and untransplanted controls for each genotype group (N = 4-6 hippocampi per condition) for scRNA-seq. **(B)** Uniform manifold approximation and projection (UMAP) clustering of isolated hippocampal microglia. 16 microglial clusters were identified. **(C–D)** Feature plots displaying strong expression of microglial markers *Cx3cr1* and *Csf1r* demonstrate that nearly all cells sequenced are of microglial identity. **(E)** Feature plot displaying expression of *Cd74* in select clusters. **(F-J)** Feature plots displaying very sparse or non-existent expression of non-microglial cell-type markers, including Syn1 (neurons), Slc17a7 (excitatory neurons), Gad1 (inhibitory neurons), Gfap (astrocytes), and Mbp (oligodendrocytes).

### Human neuronal APOE reduces homeostatic microglia in chimeric mouse hippocampus

Further analysis of the 16 cell clusters revealed several distinct clusters of interest, specifically clusters 1, 4, and 12. The percentage of cluster 1 microglia (Figures 7A–7E, green) in hEKO-E4KI mice (37.6%) was nearly double that found in hE4-E4KI mice (21.9%) or hE3-E3KI mice (23.6%) (Figure 7F). Analysis of the differentially expressed genes (DEGs) revealed upregulation of homeostatic microglial markers, including *P2ry12, Cx3cr1, Fcrls, Cst3, Crybb1, and Sparc*^50–53^, relative to other microglial clusters (Figures 7G and 7J). Additionally, cluster 1 showed strongly downregulated expression of genes associated with pro-inflammatory or activated microglia, like *Cd52, APOE, Lyz2,* and *Cd74*^54^ (Figure 7G). Based on these transcriptomic features, cluster 1 represents a homeostatic microglial subpopulation which was reduced in hE4-E4KI and hE3-E3KI mice compared to hEKO-E4KI mice, suggesting that human neuronal APOE prompts reduction of homeostatic microglia.

**Figure 7.**
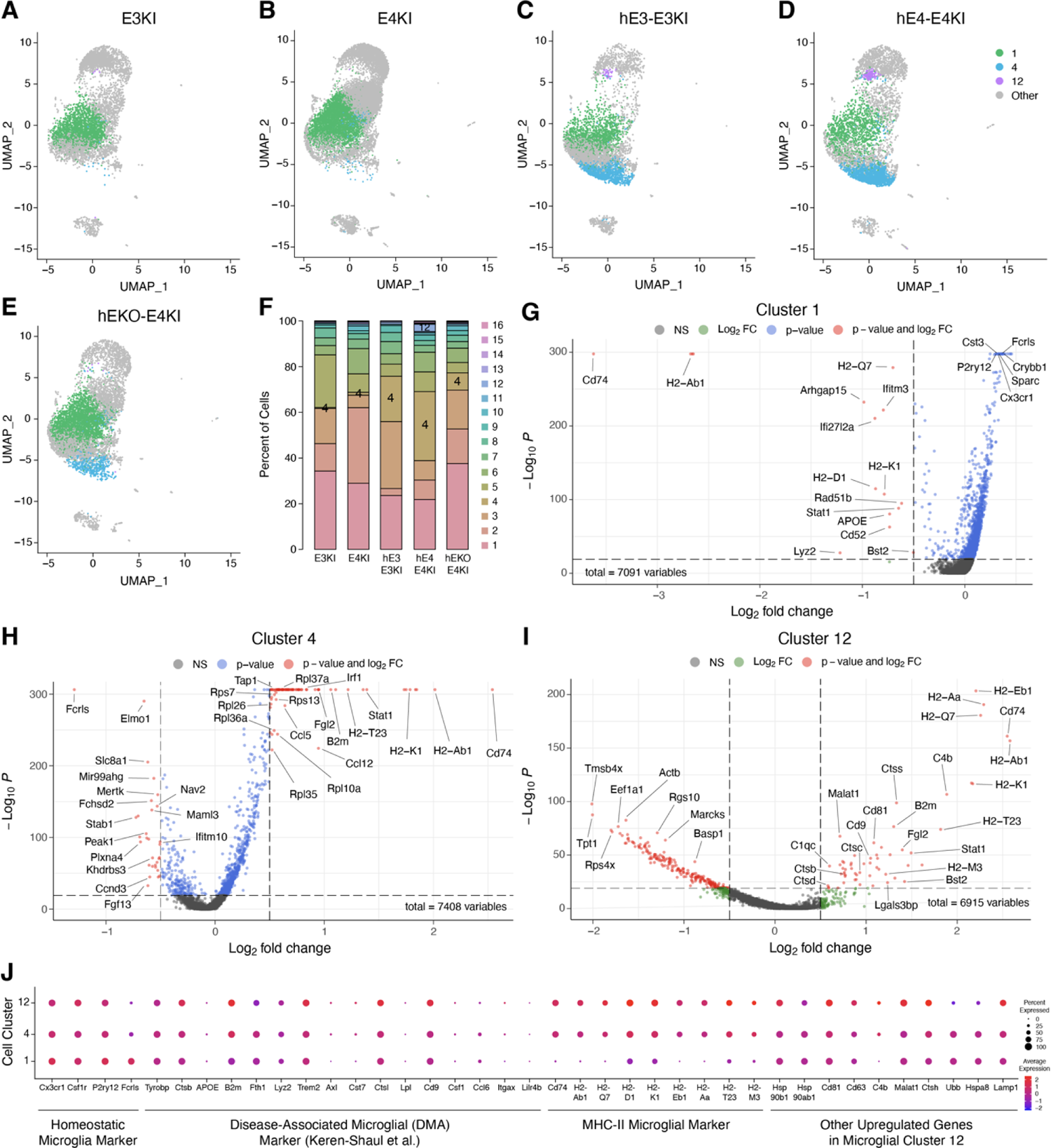
Pro-inflammatory microglia subpopulations enriched in hE4-E4KI mouse hippocampus. **(A–E)** Feature plots highlighting cells in microglial clusters 1 (green), 4 (blue), and 12 (purple). **(F)** Quantification of fraction of cells per condition for clusters 1, 4, and 12. **(G–I)** Volcano plot of the DEGs between cluster 1 (G), cluster 4 (H), or cluster 12 (I) and all other clusters. Dashed lines represent log_2_ fold change threshold of 0.5 and p-value threshold of 10 x 10^-20^. NS, not significant. **(J)** Dot plot of normalized average expression of selected marker genes for clusters 1, 4, and 12. The size of the dots is proportional to the percentage of cells expressing a given gene.

Diving further into the effect of APOE isoforms on this homeostatic microglia cluster, we examined the DEGs in hE4-E4KI microglial cluster 1 versus hE3-E3KI microglial cluster 1. This comparison showed that the hE4-E4KI microglia cluster 1 expressed increased levels of pro-inflammatory and AD-associated genes like *Cdk8* (a promoter of NF-κB signaling and chemokine expression)^55,56^, *Cmss1* (previously found to be upregulated in APOE4 male mice)^57^, *Marcks* (associated with cell motility and phagocytosis and highly expressed by senile amyloid plaque-associated microglia)^58,59^, and *Tmem176b* (expressed highly by amyloid plaque-adjacent microglia and associated with lysosomal degradation)^60^ (Figure S7A). Though we cannot determine whether these effects are attributed to microglial or neuronal APOE4, it is clear that APOE4 induces transcriptomic changes toward pro-inflammatory signatures even in homeostatic microglia compared to APOE3.

To isolate the effects of neuronal APOE4 on E4KI microglia, we examined DEGs in hE4-E4KI microglial cluster 1 versus hEKO-E4KI microglial cluster 1. We found that hE4-E4KI microglia cluster 1 showed increased expression of antigen-presentation markers (*Cd74* and *H2-K1*), vesicle/endosomal trafficking markers (*Dynll1* and *Washc4*), and microglial chemotaxis marker *P4ha1*^61–63^ (Figure S7B). Taken together, these indicate that the presence of neuronal APOE4 may prime even homeostatic E4KI microglia to assume a relatively pro-inflammatory, reactive state.

### Pro-inflammatory microglia subpopulations are more abundant in hE4-E4KI chimeric mice

In contrast to cluster 1, the percentage of microglia in cluster 4 (Figure 7A–7E, blue) was much higher in hE4-E4KI mice (30.2%) and hE3-E3KI mice (19.9%) compared to hEKO-E4KI mice (7.6%) (Figure 7F). The difference in cluster 4 microglia representation was even more stark between transplanted and untransplanted mice, with untransplanted E3KI and E4KI mice having only 0.4% and 1.3% of microglia attributed to cluster 4 (Figure 7F). These data imply that cluster 4 may represent a microglia subpopulation responding specifically to transplanted human neurons expressing APOE, especially APOE4. Examining the DEGs in this cluster more closely revealed significant upregulation of MHC-II antigen presentation genes (*Cd74, H2-Ab1, Tap1*)^64,65^ and genes involved in chemokine/interferon response signaling (*Stat1, Irf1, B2m, Ccl5, Ccl12*)^66–68^ (Figure 7H). Each of these gene subsets serves as distinct markers for MHC-II expressing (MHC-II) microglia and interferon-responsive (IFN-R) microglia, two activated microglia subpopulations previously described in AD mouse models^64^. Interestingly, cluster 4 microglia from these chimeric mice do not fully match either those MHC-II or IFN-R microglia subpopulations, instead displaying a gene expression profile with parts of both. Cluster 4 microglia also showed significant downregulation of homeostatic microglia markers (*Fcrls*, *Mertk*, *Elmo1*, *Nav2*), a signature of activated microglia. Additionally, several downregulated genes from cluster 4 matched those downregulated specifically in reactive hippocampal microglia (*Plxna4*, *Maml3*, *Fchsd2*, *Slc8a1*)^69^.

Compared to hE3-E3KI microglia, hE4-E4KI microglia from cluster 4 upregulated a variety of pro-inflammatory markers, once again including genes from MHC-II microglia (*H2-Ab1, H2-Aa, Ly6e*) and IFN-R microglia (*Stat1*) expression profiles (Figure S7C). Cluster 4 microglia from hE4-E4KI mice also showed increased levels of *Cdk8* and *Malat1*, both potent activators of the microglia inflammasome^55,56,70,71^. Comparison of hE4-E4KI versus hEKO-E4KI microglia for cluster 4 also yielded increased levels of MHC-II microglia genes (*H2-Aa, H2-Eb1*), along with additional pro-inflammatory markers (*Cd74, Stat1, Irf1)* (Figure S7D). *Lamp1* and *Lyz2*, both lysosomal genes that serve as markers of activated phagocytic microglia^72^, were also upregulated in hE4-E4KI microglia, indicating that human neuronal APOE4 may promote a microglial state conducive to increased uptake/engulfment. When we used Gene Ontology (GO) analysis to investigate differential regulation of pathways in hE4-E4KI compared to hEKO-E4KI microglia, we found that hE4-E4KI cluster 4 microglia showed alterations in pathways related to MHC protein complex, antigen processing/presentation, cellular response to interferons, and cytokine response (Figure S7E). Interestingly, cytokine response and other pro-inflammatory genes were also upregulated in *in vivo* human microglia from a chimeric mouse model in response to amyloid^73^, suggesting that the pro-inflammatory response in our chimeric mice is also relevant to the response of human microglia in AD. These data, taken together, indicate that human neuronal APOE, especially APOE4, induces pro-inflammatory microglial subpopulations, such as MHC-II microglia.

Microglial cluster 12 (Figures 7A–7E, purple), although a relatively small cluster, was highly represented in hE4-E4KI mice. The percentage of microglia in hE4-E4KI mice was 3.2% (Figure 7F), over 3-fold higher than the percentage in hE3-E3KI mice (0.9%) and over 30-fold higher than the percentage in hEKO-E4KI mice (0.09%) (Figure 7H). In fact, hEKO-E4KI mice have a similar low percentage of microglia in cluster 12 as untransplanted mice (E3KI: 0.1%, E4KI: 0.01%) (Figure 7F). This marked difference in the proportion of cluster 12 microglia between hE4-E4KI and hEKO-E4KI mice may indicate that cluster 12 represents a microglial subpopulation that responds more strongly to transplanted human neurons with APOE, particularly APOE4. A closer look at the DEGs in microglial cluster 12 showed even more pro-inflammatory genes upregulated than in microglial cluster 4. Although cluster 12 microglia did show upregulated levels of some markers associated with disease-associated microglia (DAM), including *Trem2, Ctsl, and Cd9*, their expression profile does not fully match the DAM profile previously outlined^74^. Instead, the gene expression profile of cluster 12 microglia most closely resembles MHC-II microglia, with a very high proportion of upregulated DEGs falling into the MHC antigen presentation pathway: *Cd74, H2-Ab1, H2-Aa, H2-Eb1, H2-K1, and B2m*^64,75^ (Figure 7I, 7J). Another study showed that MHC-II microglia exhibit increased Aβ phagocytic capacity than other microglia in APP/PS1 AD model mice^76^. In line with these findings, cluster 12 microglia also exhibit a number of upregulated lysosomal genes (*Lgals3bp, Ctsb, Ctsc, Ctss, Ctsd)* associated with increased microglial phagocytosis and AD risk^77,78^. Additionally, microglial exosome genes *Cd9* and *Cd81* were highly upregulated in cluster 12 microglia. As exosomes have previously been identified as a potential mechanism for pathological amyloid and tau deposition^79,80^, it is interesting to find significant upregulation of exosome markers in a microglial cluster particularly abundant in the hE4-E4KI mice, which had high levels of amyloid and tau pathologies.

Perhaps equally interesting is the cell expression pattern of *Cd68*, which is a lysosomal protein that is highly expressed in activated phagocytic microglia in AD models^81–83^. Several studies have also shown that expression of *Cd68*, unlike several other microglial activation markers, was consistently increased in post-mortem brains with AD and APOE4 relative to controls^84^ and was associated with neurodegeneration and cognitive decline in aging^20,85^. When we examined our scRNA-seq dataset, *Cd68* expression was generally higher in most clusters for E4KI microglia compared to E3KI microglia (Figure S7F). However, clusters 4 and 12 of hE4-E4KI microglia showed the highest *Cd68* expression compared to any other cluster or chimeric condition (Figure S7F). Fittingly, as mentioned earlier, clusters 4 and 12 display upregulated markers of MHC-II microglia, which also highly express *Cd68* in inflammatory AD model mice^76^. Together, these data support the conclusion that microglial clusters 4 and 12, both of which are enriched in hE4-E4KI mice, are highly reactive and pro-inflammatory microglia subpopulations.

## DISCUSSION

In this study, we examine the effects of microglial presence and neuronal APOE on AD pathology in an *in vivo* chimeric disease model. First, we demonstrated that the APOE4 isoform increased the number of Aβ, Thioflavin-S, and p-tau aggregates in hE4-E4KI mice compared to hE3-E3KI mice. These results match a litany of previous studies, including our previous chimeric AD model study, that show APOE4’s exacerbation of amyloid and/or tau pathology^12,19,21,25,37,45^. We also explored the less well-established effect of microglia depletion on human neuronal APOE4-driven AD pathologies. Past studies of microglial depletion in AD mouse models presented mixed results: microglia depletion reduced Aβ plaque burden in 5XFAD mice^34,35,86,87^, reduced tau pathology in PS19/P301S tauopathy mice^20,79^, had no effect on pathology in aged APP/PS1 or 5XFAD or 3xTg-AD tau mice^40,88–90^, and even increased tau spread in tau-injected 5XFAD mice^91^. Our results demonstrated that, in an AD model with human neurons in an *in vivo* context with no early-onset AD mutations, depletion of microglia reduced Aβ and tau aggregates in hE4-E4KI mice, indicating that microglia promote human neuronal APOE4-driven Aβ and tau pathologies. As discussed in our previous paper establishing this model, the formation of aggregates in our chimeric disease mice have a particular relevance to human late-onset AD^25^, signifying strong potential for translatability of these findings.

Similar to our results in hEKO-E4KI mice, a recent study in APP/PS1 AD model mice demonstrated that general removal of APOE induced more diffuse morphology of amyloid aggregates^92^. Importantly, our results identify neurons as a relevant source of APOE for compaction of amyloid aggregates. Interestingly, pathological analyses in post-mortem human brains showed that diffuse amyloid plaques tend to appear in the brains of older cognitively normal patients rather than AD patients, who tended to have more dense-core plaques^93^. These results suggest that the diffuse amyloid deposits formed in hEKO-E4KI mice may not be pathologically detrimental, a finding that aligns with studies showing that removal of APOE, especially neuronal APOE4, is mostly associated with amelioration of AD-related neurodegeneration and cognitive deficits in AD mouse models^21,37,94,95^.

The significant increase in transplant-localized APOE upon microglial depletion in our chimeric model bears a striking resemblance to a previously reported finding that microglial depletion in PS19-E4 tauopathy mice led to strongly increased APOE in neurons in the hippocampus^20^. Both studies suggest that microglia normally clear APOE produced by neurons; thus, microglial depletion prevents this clearance and leaves behind the aforementioned APOE from neurons. Since both Aβ and p-tau pathologies were reduced upon microglia depletion in hE4-E4KI mice, together these data support a hypothesis that neuronal APOE4-driven Aβ and p-tau pathologies require microglial participation.

Finally, examination of the gene expression profile of hippocampal microglia in the chimeric mice revealed three main microglial clusters of interest: homeostatic microglial cluster 1 and pro-inflammatory microglial clusters 4 and 12. The gene expression profiles of clusters 4 and 12 partially resembled AD-related microglial subpopulations IFN-R and DAM, but the highest profile alignment for both clusters (particularly cluster 12) was with MHC-II microglia. Though less well-studied than DAM, microglia that specifically express high levels of MHC-II markers have been shown to be enriched in human AD patient brains^96,97^. Past studies have also noted that, although MHC-II microglia often express subsets of DAM markers^98^, MHC-II microglia—not DAM—were specifically associated with increased neuron loss and neuropathology in neurodegeneration model mice^75,99^. As these MHC-II microglia in clusters 4 and 12 are particularly found in hE4-E4KI mice, it is possible that these cells are involved in the higher AD pathology found in hE4-E4KI mice. The relevant mechanisms by which MHC-II microglia may contribute to pathogenesis are as yet unknown. While MHC-II microglia are primarily known for their antigen presentation functions, they have also displayed upregulation of Aβ phagocytic capacity in APP/PS1 transgenic mice and may have other currently unidentified roles in the brain^76,100^. In addition, the relationship between clusters 4 and 12 remain unclear. They may represent two separate subpopulations of microglia reacting to different stimuli, with cluster 4 more generally responding to transplanted human neurons and cluster 12 more specifically responding to effects driven by neuronal APOE, especially APOE4. Alternatively, it is possible that cluster 4 microglia, which displays a tempered upregulation of many of the same DEGs as cluster 12, may represent an intermediate state of microglia between the more homeostatic cluster 1 microglia and the more inflammatory MHC-II microglia in cluster 12. Further studies are necessary to distinguish these two possibilities.

APOE4 generally increased pro-inflammatory markers in most microglia clusters of interest, including homeostatic cluster 1, and neuronal APOE4 in particular tended to upregulate pathways related to antigen presentation and interferon response in both cluster 4 and cluster 12 microglia. Interestingly, the specific combination of pro-inflammatory microglial clusters 4 and 12 and neuronal APOE4 in hE4-E4KI mice resulted in dramatically increased expression of AD pathology-associated lysosomal gene *Cd68* in those cells, suggesting that they are highly active and more inflammatory. Interestingly, a recent study found that APOE4 microglia showed a reduced pro-inflammatory response (compared to APOE3 microglia) in an *ApoE-KO* amyloid mouse model^101^, and another paper demonstrated that deleting APOE4 from microglia in APP/PS1:*APOE4-KI* mice restored microglial reactivity^102^. In light of these findings, our results indicate that the upregulation of pro-inflammatory markers in hE4-E4KI microglia is potentially a microglial response specific to human neurons, particularly those expressing APOE4, or to Aβ and p-tau protein without pathological mutations. Depletion of specific pro-inflammatory clusters may be important for reduction of AD-related pathologies in hE4-E4KI-PLX mice.

### Limitations of Study

There are limitations to consider in this study. As previously mentioned, although microglia were significantly depleted in all conditions, depletion in hE3-E3KI-PLX mice was less effective than in hE4-E4KI-PLX mice. Accordingly, the lack of effect of microglial depletion on Aβ and p-tau pathology in hE3-E3KI-PLX mice may indicate either that APOE3 microglia play a limited role in pathology formation or that depletion of key microglial populations in these mice was insufficient to produce a significant effect. Additionally, while our studies explore the role of mouse microglia in AD pathogenesis, it is important to validate the translatability of these and other observations with human microglia. Of particular interest is a previously published chimeric model engrafting human stem cell-derived microglia into the mouse brain^73^. A combined chimeric mouse model with both human neurons and microglia could serve as a powerful tool to explore human-specific AD pathogenesis. Finally, it is unclear whether the formation of Aβ and p-tau aggregates worsens neurotoxicity^87,103,104,105,106^ or serves as a protective process to limit the effects of pathological proteins^107–109^. One recent study in an APOE4 tauopathy mouse model demonstrated that microglial presence promotes both tau pathology and neurodegeneration^37^, suggesting that microglial removal may be therapeutically beneficial. However, further confirmation of whether depleting microglia and reducing aggregates will ameliorate or exacerbate neurodegeneration and cognitive deficits in AD is essential before pursuing microglial depletion as a possible therapeutic avenue.

## Acknowledgments

This work was partially supported by the National Institutes of Health grants R01AG071697, R01AG076647, 1R01AG065540, and P01AG073082 to Y.Huang, F31AG074672 to N.K., and F31AG074690 to M.R.N. This work was also supported by the National Science Foundation Graduate Research Fellowship to A.R. under Grant No. 2034836. We thank the Huang Lab staff for their valuable discussions about the experimental design as well as data analyses and interpretation. We also thank Dr. Wenjie Mao and Dr. Andrew Mendiola for guidance on effective microglia collection and isolation. Some figures were partially created in BioRender.com. Gladstone Institutes Flow Cytometry Core was supported by a NIH grant S10 RR028962 and the James B. Pendleton Charitable Trust for use of the FACSAria II and Fortessa X-20, DARPA for the use of Fortessa X-20, and NIH P30 AI027763 for use of the FACSAria II.

## Author Contributions

A.R. and Y.Huang designed and coordinated the studies and wrote the manuscript. A.R. performed the majority of studies and data analyses. M.J.K., O.Y., J.B., M.N., Z.L., and N.K. helped with some immunohistochemical studies, data collection, and sample preparation for scRNA-seq analyses. J.B. helped with most mouse brain collections. Y.Hao isolated cell nuclei and prepared samples for scRNA-seq. N.C. performed the majority of the scRNA-seq analyses, with help from B.G. and L.D. S.Y.Y. and P.A. managed all mouse lines and helped with some mouse brain collections. Y.Huang supervised the project.

---

**Figure S1.**
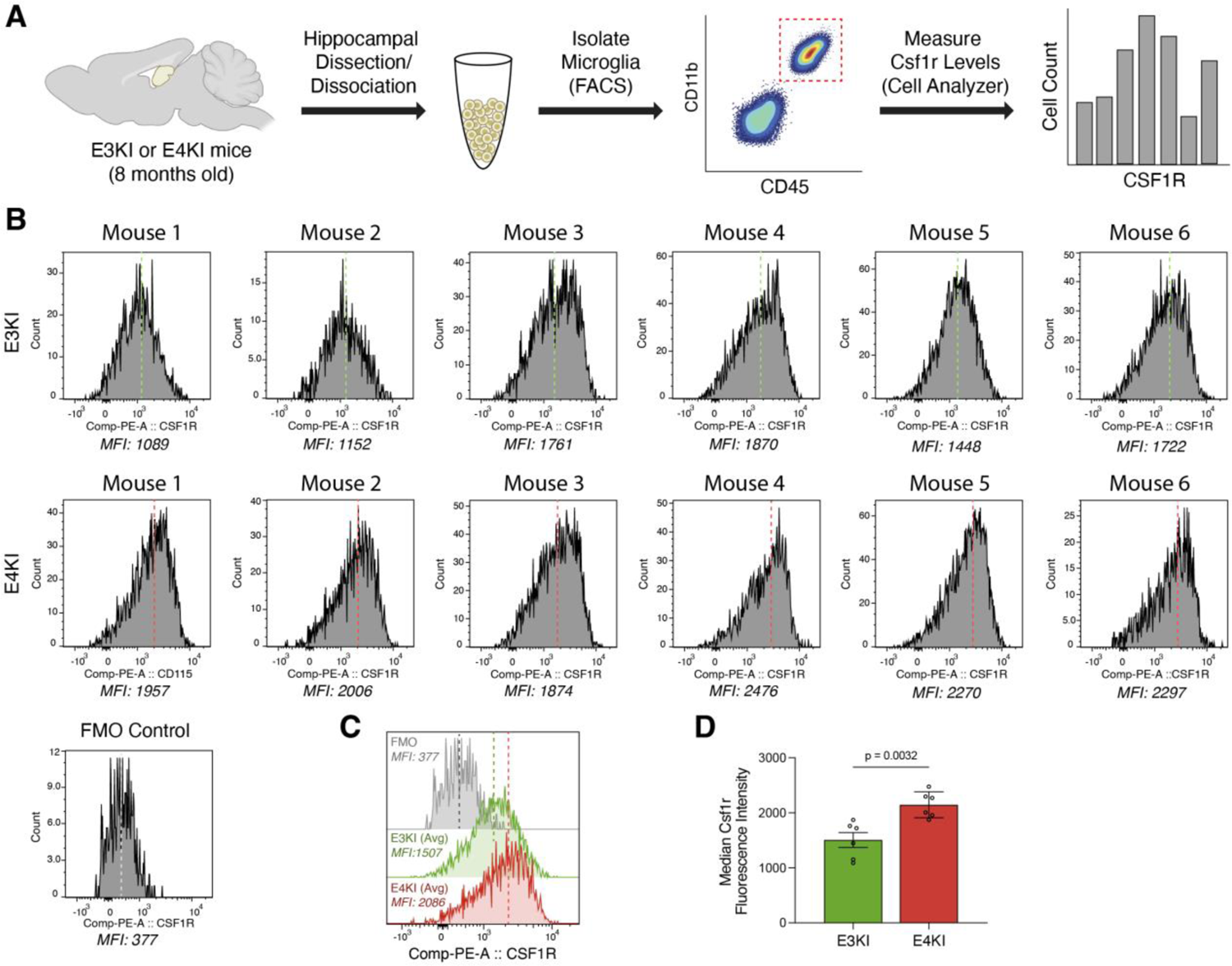
E4KI mouse microglia express higher levels of Csf1r than E3KI mouse microglia, related to Figure 2. **(A)** Workflow summarizing isolation and FACS sorting of CD11b^+^/CD45^int^ microglia from untransplanted E3KI and E4KI mouse hippocampi, and measurement of Csf1r levels by cell analyzer. **(B)** Histograms of fluorescence intensity of Csf1r PE-conjugated antibody in individual E3KI and E4KI mice and a fluorescence minus one (FMO) control. Dotted line represents median fluorescence intensity (MFI) of each sample. **(C)** Histogram overlaying Csf1r expression levels for the FMO control and a representative mouse each for E3KI and E4KI conditions. Dotted line represents average MFI for FMO and each condition. **(D)** Quantification of MFI for hippocampal microglia from untransplanted E3KI and E4KI mice. Each dot represents one mouse per condition (E3KI, n=6; E4KI, n=6). All data are expressed as mean ± S.E.M. Differences between groups were determined by unpaired two-tailed t-test.

**Figure S2.**
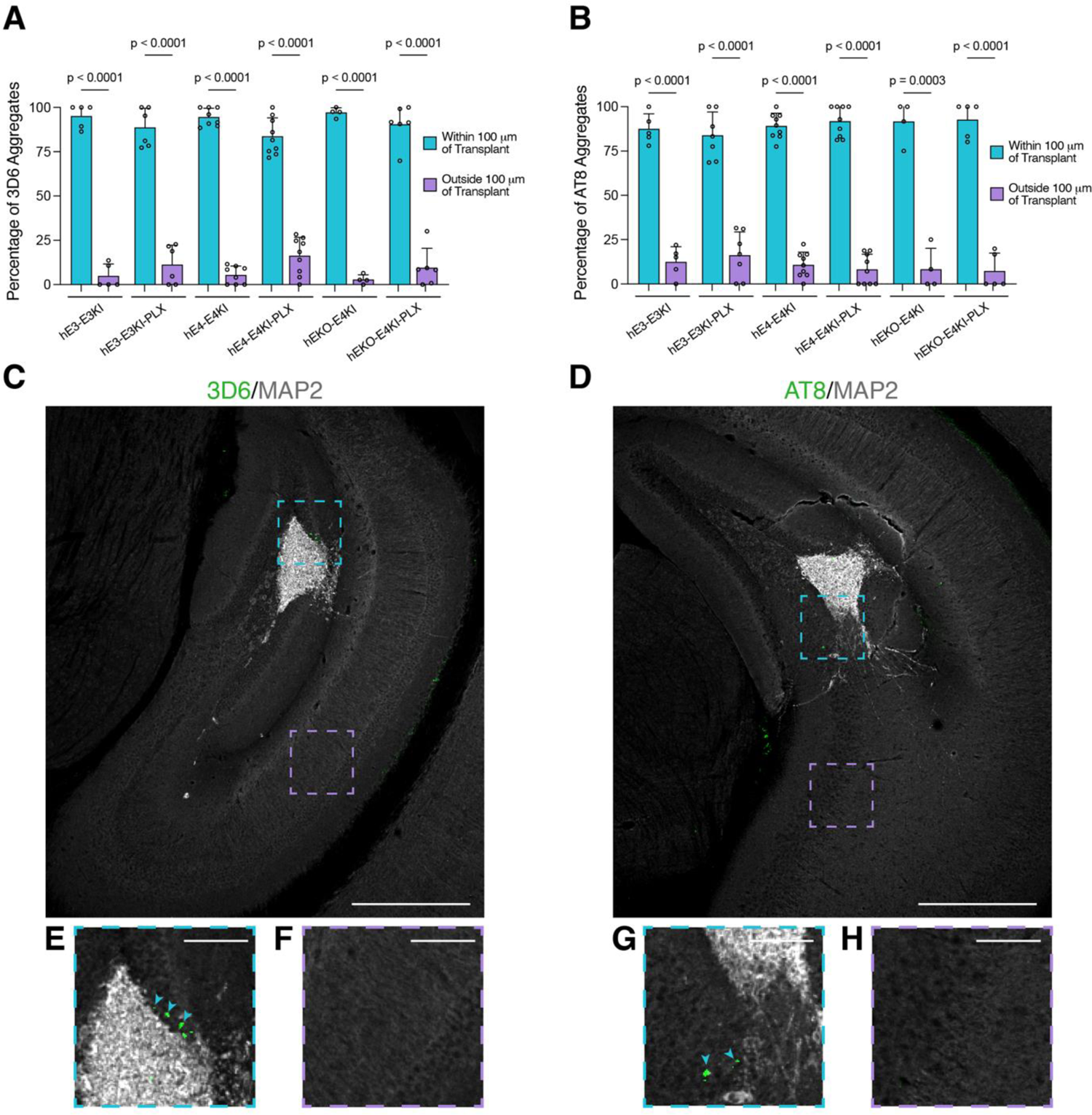
Aβ and p-tau aggregates form primarily within 100 μm of human neuron transplants, related to Figures 3 and 4. (**A** and **B)** Quantification of percentage of hippocampal 3D6^+^ Aβ aggregates (hE3-E3KI, n=5; hE3-E3KI-PLX, n=6; hE4-E4KI, n=8; hE4-E4KI-PLX, n=9; hEKO-E4KI, n=4; hEKO-E4KI-PLX, n=6) (A) or AT8^+^ p-tau aggregates (hE3-E3KI, n=5; hE3-E3KI-PLX, n=7; hE4-E4KI, n=9; hE4-E4KI-PLX, n=9; hEKO-E4KI, n=4; hEKO-E4KI-PLX, n=5) (B) within or outside 100 μm of human neuron transplants. Each dot represents one mouse per condition. All data are expressed as mean ± S.E.M. Differences between groups were determined by two-way ANOVA with Benjamini’s post hoc test for multiple comparisons. **(C** and **D)** Representative immunohistochemical images of hippocampal distribution of 3D6^+^ Aβ aggregates (C) or AT8^+^ p-tau aggregates (B) relative to MAP2^+^ human neuronal transplants (gray). Scale bar, 500 μm. **(E** and **F)** Magnified insets of C, highlighting the presence (E) or absence (F) of 3D6^+^ Aβ aggregates within or outside 100 μm of the human neuronal transplants. Scale bar for insets, 100 μm. (**G** and **H**) Magnified insets of D, highlighting the presence (G) or lack (H) of AT8^+^ p-tau aggregates within or outside 100 μm of the human neuronal transplants. Scale bar for insets, 100 μm.

**Figure S3.**
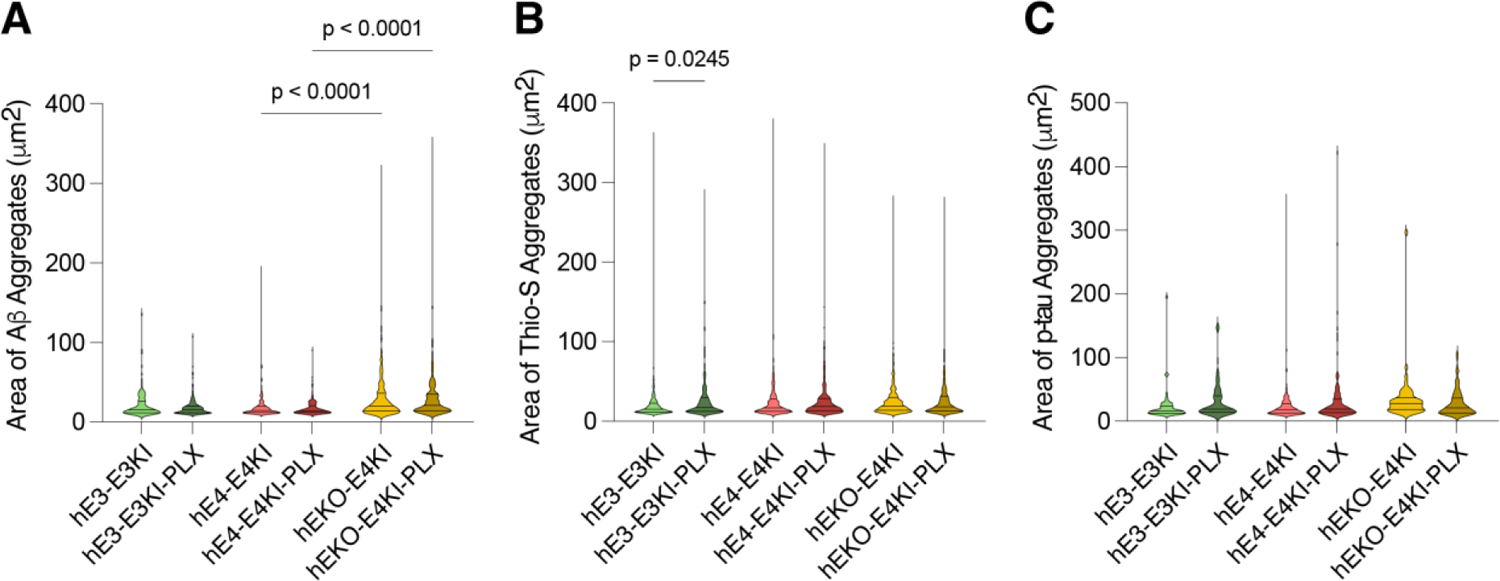
Aβ and p-tau aggregate sizes across conditions, related to Figures 3 and 4. **(A–C)** Violin plot quantification of aggregate size across conditions for 3D6^+^ Aβ aggregates (A), Thioflavin-S^+^ dense-core Aβ aggregates (B), and AT8^+^ p-tau aggregates (C). Differences between groups were determined by two-way ANOVA with Benjamini’s post hoc test for multiple comparisons.

**Figure S4.**
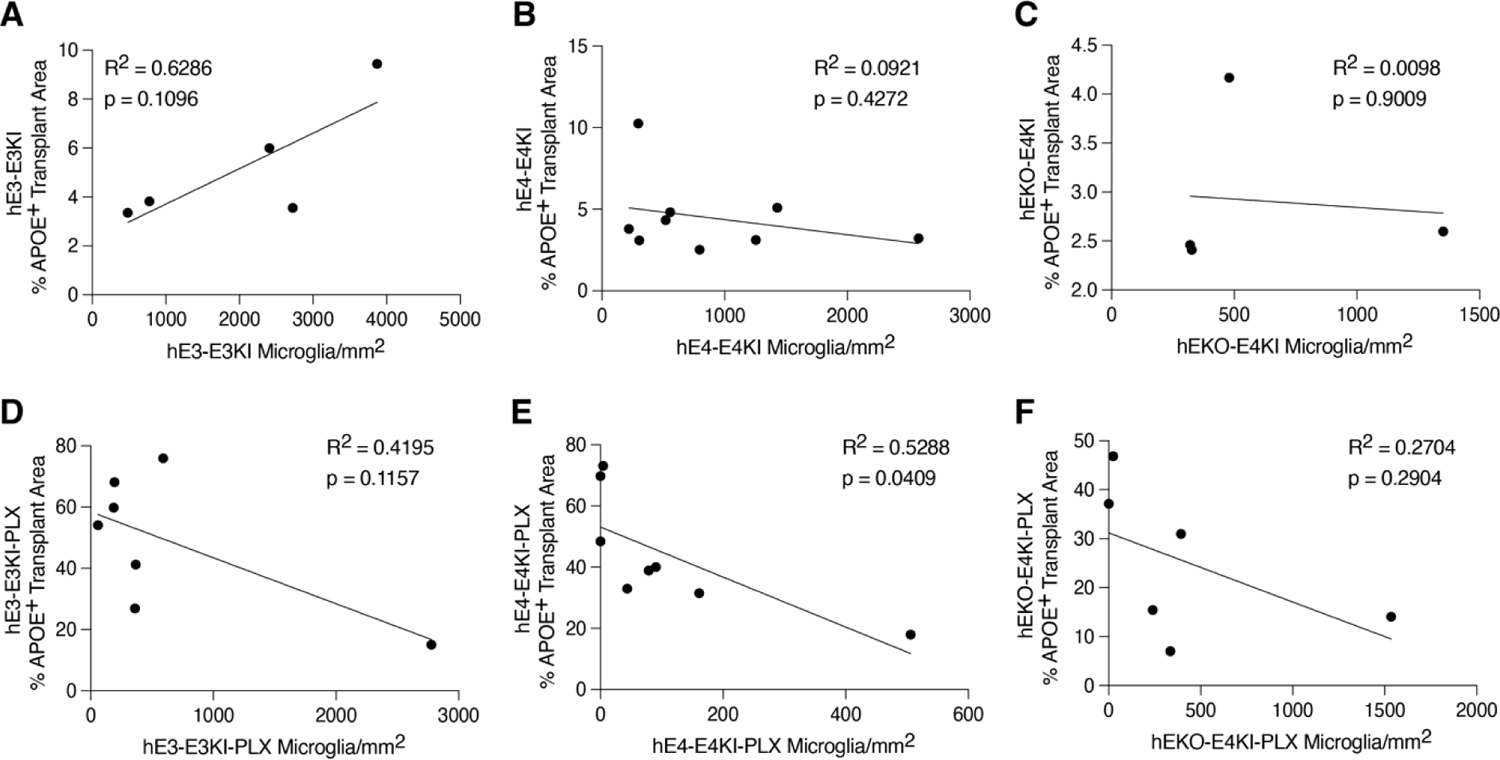
APOE levels in human neuron transplants correlates negatively with microglia density in hE4-E4KI-PLX mice, related to Figure 5. **(A–F)** Correlations for each condition between microglia density and APOE^+^ area coverage within human neuronal transplants. Each dot represents one mouse per condition (hE3-E3KI, n=5; hE3-E3KI-PLX, n=7; hE4-E4KI, n=9; hE4-E4KI-PLX, n=8; hEKO-E4KI, n=4; hEKO-E4KI-PLX, n=6). Pearson’s correlation analyses (two-sided).

**Figure S5.**
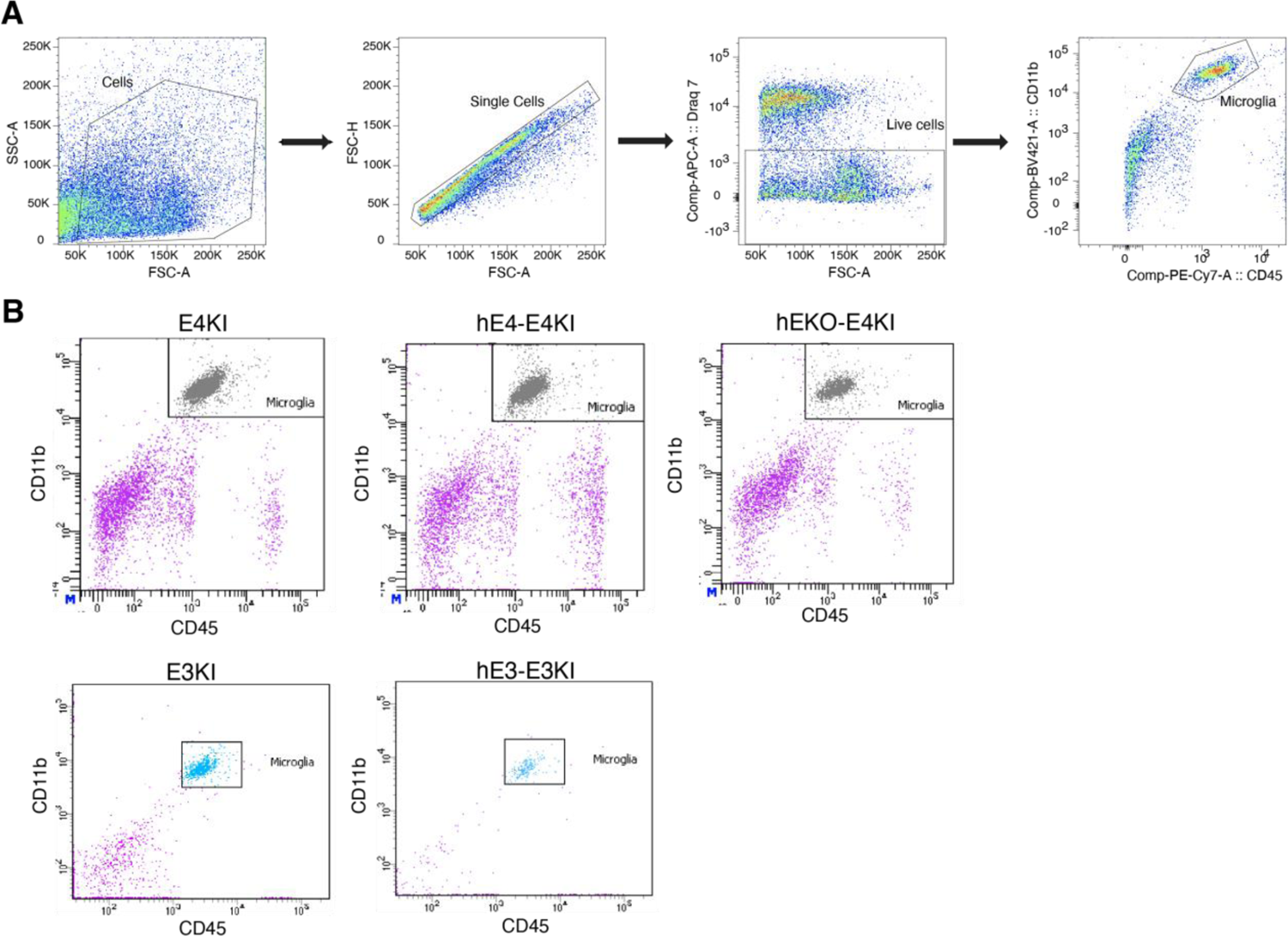
FACS gates to isolate microglia for scRNA-seq, related to Figures 6 and 7. **(A)** Representative flow cytometry gating strategy using CD11b and CD45 antibodies to isolate microglia for scRNA-seq. **(B)** Flow cytometry gates for final microglia isolation step for each chimeric mouse condition.

**Figure S6.**
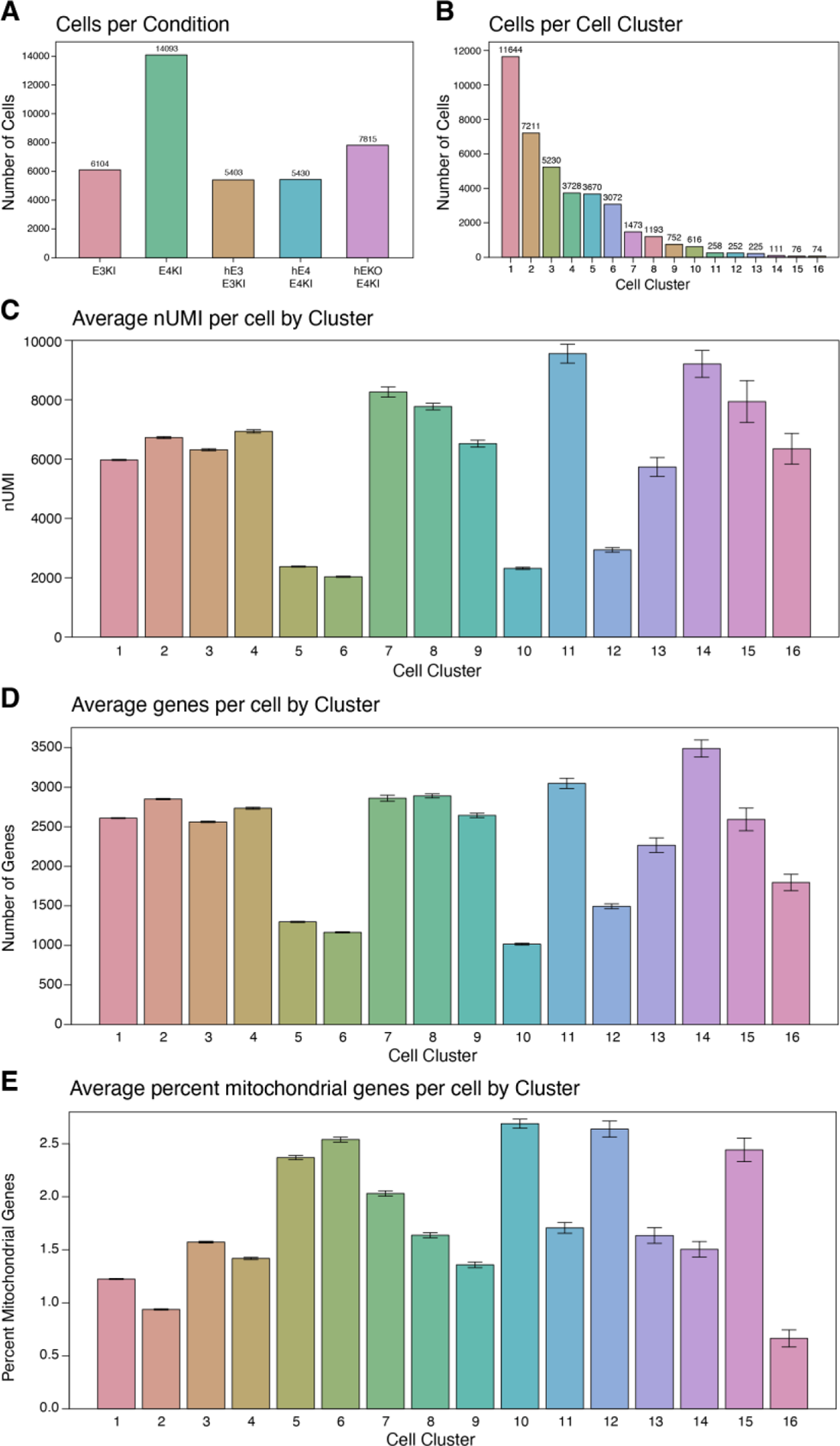
Quality control measures in scRNA-seq analysis of chimeric mouse hippocampal microglia, related to Figures 6 and 7. **(A)** Quantification of number of cells sequenced per cluster. **(B)** Quantification of number of cells sequenced per condition. **(C)** Quantification of the number of genes per cell in each cluster. Data are expressed as mean ± S.E.M. **(D)** Quantification of nUMI per cell for each cluster. **(E)** Quantification of the percentage of mitochondrial genes per cell in each cluster. Data are expressed as mean ± S.E.M.

**Figure S7.**
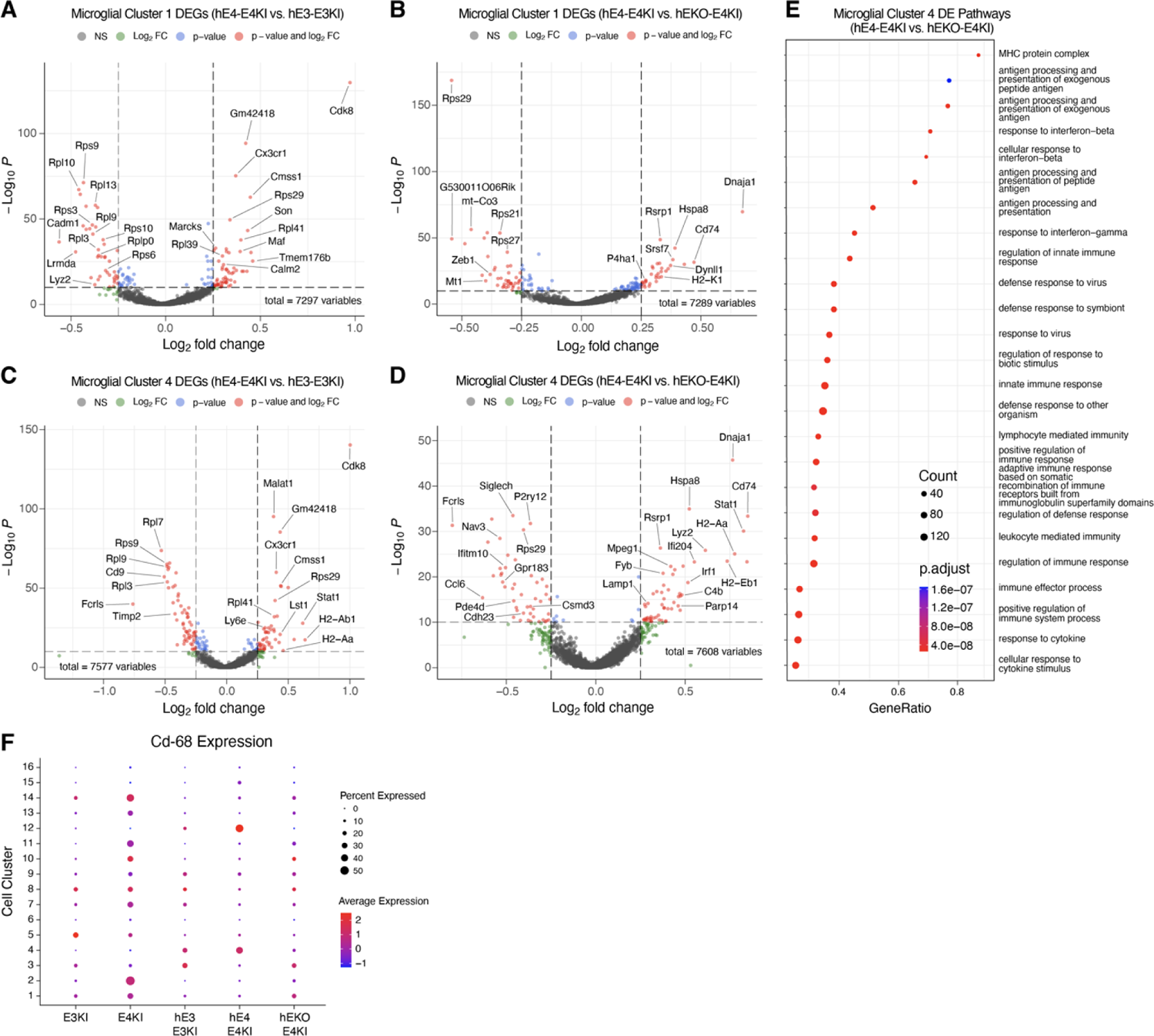
Differential responses of E3KI versus E4KI microglia to respective human neuron transplants and of E4KI microglia to E4 versus EKO human neuron transplants, related to Figure 7. **(A** and **B)** Volcano plots of the DEGs for hippocampal microglia cluster 1 in hE4-E4KI versus hE3-E3KI mice (A) and in hE4-E4KI versus hEKO-E4KI mice (B). Dashed lines represent log_2_ fold change threshold of 0.25 and p-value threshold of 10 x 10^-10^. NS, not significant. **(C** and **D)** Volcano plots of the DEGs for hippocampal microglia cluster 4 in hE4-E4KI versus hE3-E3KI mice (C) and in hE4-E4KI vs hEKO-E4KI mice (D). Dashed lines represent log_2_ fold change threshold of 0.25 and p-value threshold of 10 x 10^-10^. NS, not significant. **(E)** GO dot plot of DE pathways for hippocampal microglia cluster 4 in hE4-E4KI versus hEKO-E4KI mice. The size of the dots is proportional to the number of DE genes within that pathway. **(F)** Dot plot of normalized average gene expression of *Cd68* for all clusters in each condition.

## Supplementary Tables

**Table S1. ScRNA-seq data related to** Figure 7**, including DEGs in clusters 1, 4, and 12 versus other clusters and percent of cells per condition assigned to each cluster.**

**Table S2. ScRNA-seq data related to Figure S6, including cells per condition, cell per cell cluster, nUMI per cell by cluster, genes per cell by cluster, and percent mitochondria per cell by cluster.**

**Table S3. ScRNA-seq data related to Figure S7, including DEGs and pathways in hE4-E4KI vs hE3-E3KI and hE4-E4KI vs hEKO-E4KI for select clusters.**

## STAR METHODS

### Mice

Mice with human APOE3 or APOE4 knocked-in at the mouse *Apoe* locus on a C57BL/6 background were originally obtained from Taconic^110^. All animals were bred in-house using trio breeding producing 8– 10 pups per litter on average, which were weaned at 28 days. Male littermates at 3-5 months of age were randomly assigned to experimental groups. Animals were housed in a pathogen-free barrier facility on a 12 hr light cycle (lights on at 7 am and off at 7 pm) at 19–23°C and 30%–70% humidity. Animals were identified by ear punch under brief isofluorane anesthesia and genotyped by polymerase chain reaction (PCR) of a tail clipping at weaning. All animals otherwise received no procedures except those reported in this study. All animal experiments were conducted in accordance with the guidelines and regulations of the National Institutes of Health, the University of California, and the Gladstone Institutes under IACUC protocol AN117112.

### Cell Lines

All hiPSC lines were derived from human skin fibroblasts from donors and reprogrammed as previously described ^12^ and were maintained at 37°C with 5% humidity. All hiPSC lines were characterized for normal pluripotency gene expression, apoE genotypes, karyotypes, and capability of differentiating into neural stem cells as well as different types of neurons in culture. All hiPSC lines were tested negative for mycoplasma.

### hiPSC Culture

The E4/4 hiPSC line was generated as described^111,112^ from skin fibroblasts of a subject with an APOE4/4 (E4) genotype. The isogenic E3/3 (E3) hiPSC line was generated from this parental E4/4 hiPSC line as previously described^12^. hiPSCs were maintained in mTeSR medium (85850, StemCell Tech) on 6-well plates precoated with hESQ, LDEV-free Matrigel (354277, Corning). The medium was changed daily, and cells were routinely passaged 1:10–1:15 using Accutase (NC9464543, Fishersci) for dissociation. Rho kinase (ROCK) inhibitor (1254, Tocris) was added to medium at 10µM on day of passaging.

### Neuronal Differentiation of hiPSCs

hiPSCs were differentiated into neurons as previously described^12,25^, with slight modifications to increase yield. Briefly: hiPSCs were dissociated with Accutase for 5–8 minutes before being quenched with warm (37°C) N2B27 medium made of 1:1 DMEM/F12 (11330032, Thermo Fisher) and Neurobasal Media (21103049, Thermo Fisher), 1% N2 Supplement (21103049, Thermo Fisher), 1% B27 (17504044, Thermo Fisher), 1% MEM Non-essential Amino Acids (11140050, Thermo Fisher), 1% Glutamax (35050061, Thermo Fisher), and 0.5% penicillin–streptomycin (15140122, Thermo Fisher). Dissociated hiPSCs were pelleted by centrifugation, resuspended in embryoid body media [10µM SB431542 (1614, Tocris) and 0.25µM LDN (04-0074, Stemgent) in N2B27] with 10µM ROCK inhibitor (1254, Tocris), and grown in suspension in a T-75 flask (12-565-349, Fisher Scientific). Flasks were shaken manually once per hour for the first 3 hrs of incubation. On day 2, embryoid bodies had formed, and fresh embryoid body medium was replaced (embryoid bodies pelleted, old medium aspirated, cells resuspended in fresh medium) to remove ROCK inhibitor. Embryoid body medium was replaced similarly on days 4 and 6. On day 8, spheres were plated as neural progenitors onto a 10cm dish precoated with growth factor reduced (GFR) Matrigel (CB-40230A, Fisher Scientific). Neural progenitors were allowed to form neuronal rosettes and sustained in N2B27 media alone for days 8–15. Half of the media was replaced every 48–72 hrs depending on confluency and media consumption. Neuronal rosettes were lifted on day 16 using STEMdiff™ Neural Rosette Selection Reagent (05832, StemCell Tech) as directed by manufacturer and plated into 3 wells of a 6-well plated precoated with GFR Matrigel in N2B27 with 100ng/ml FGFb (100-18B, Peprotech) and 100ng/ml EGF (AF-100-15, Peprotech). This N2B27 medium with FGFb and EGF was replaced daily. On day 20, neural progenitors were dissociated with Accutase, quenched with N2B27, and resuspended in STEMdiff™ Neural Progenitor Medium (05833, StemCell Tech) at 1.2×10^6^ cells/2ml for 1 well of 6-well plate, precoated with GFR Matrigel. Neural progenitor cells were fed with fresh Neural Progenitor Medium daily. On day 28, medium was switched to complete neuronal medium (10ng/ml BDNF (450-02, Peprotech) and 10ng/ml GDNF (450-10, Peprotech) in N2B27) with 10nM DAPT (2634, Tocris). Cells were fed with fresh complete neuronal medium daily for 7 days and then harvested for cell transplantation.

### Cell Transplantation Preparation

hiPSC-derived neurons (D35-42) were washed in 1X PBS then incubated in warm Accutase (Millipore) for 15 min or until neurons dissociated with gentle tapping. Accutase (Millipore) was neutralized with N2B27 medium to bring total volume to around 30 mL and then cells were filtered through a 40 μm strainer (Fisher) to ensure a single cell suspension. Single cells were then centrifuged and resuspended to concentration of 500 cells/nL in 1X HBSS (GIBCO) supplemented with 10 ng/mL BDNF (Peprotech), 10 ng/mL GDNF (Peprotech) and 100 ng/mL DNaseI (Roche) and kept at 4°C until transplantation.

### Stereotaxic Surgery for Cell Transplantation

Mice were anesthetized with an intraperitoneal injection of ketamine (60 mg/kg) and xylazine (30 mg/kg) and maintained on 0.8%-1.0% isofluorane (Henry Schein). Mice were secured in a stereotaxic alignment system model 940 using earbars and a tooth bar (Kopf Instruments). The scalp was prepared by removing hair with scissors and sterilizing with 70% ethanol. The scalp was then cut open using a scalpel and sterilized with 70% ethanol. The cranial sutures were better visualized using 3% hydrogen peroxide. Following identification of Bregma, bilateral rostral and caudal stereotaxic sites were drilled with a 0.5 mm microburr (Fine Science Tools) using coordinates X = ± 1.65, Y = 2.00, Z = 1.7 and X = ± 2.90, Y = 3.20, Z = 2.2, with Z measured from the surface of the brain. Cell suspensions (500 cells/nL) were loaded into 60 μm tip diameter, 30° beveled glass micropipette needles (Nanoject, Drummond Scientific Company). Approximately 30,000 cells per site were injected at 4 sites (2 per hemibrain) at a rate of 25 nL/sec and allowed to diffuse for 1 min. Each mouse received four total cell transplants, two per hemibrain hippocampus. Following surgery, mice were sutured with nylon monofilament non-absorbable 6-0 sutures (Henry Schein), and administered analgesics buprenorphine (0.0375 mg/kg intraperitoneally), ketofen (5 mg/kg subcutaneously), and saline (500µL intraperitoneally). Mice were monitored on a heating pad until ambulatory and provided Hydrogel for hydration. Immunosuppressants were administered immediately after transplantation (day 0) via intraperitoneal injection followed by injections on day 2, 4, and 6 post transplantation. Immunosuppressants were comprised of a cocktail of anti-mouse CD40L (CD154) (BioXCell), anti-mouse CTLA-4 (CD152) (BioXCell), and anti-mouse LFA-1a (CD11a) (BioXcell), and all were used at a concentration of 20 mg/kg.

### PLX3397 and Control Chow

PLX3397 (Pexidartinib, HY-16749) was provided by MedChemExpress. For PLX chow, PLX3397 was incorporated into AIN-76A chow (Research Diets) at a concentration of 400mg/kg. Chimeric mice were fed either a control AIN-76A chow for 8 months, or fed control AIN-76A chow for 4 months and then PLX chow for 4 months.

### Collection of Mouse Tissue

Mice were deeply anesthetized with intraperitoneal injections of avertin (Henry Schein) and transcardially perfused for 1 min with 0.9% saline. Right hemi-brains were drop-fixed in 4% paraformaldehyde (15710, Electron Microscopy Sciences), rinsed in PBS (Corning) for 24 hrs, and cryoprotected in 30% sucrose (Sigma) for 48 hrs at 4°C. The fixed right hemi-brains were sliced into 30µm coronal sections spanning the hippocampus on a freezing sliding microtome (Leica) and stored in cryoprotectant (30% Ethylene Glycol, 30% Glycerol, 40% 1X PBS) at −20°C. For the left hemi-brains, the hippocampus was dissected out and further prepared for microglia isolation and single-cell RNA-sequencing.

### Immunohistochemistry

Multiple sections from each mouse (30µm thick, 300µm apart) were transferred to a 24-well plate and washed 2×5min with PBS to remove cryoprotectant. Sections were treated with UV radiation overnight in PBS. The next day, sections were washed 2×5min in PBS-T (PBS + 0.1% Tween-20), then incubated in PBS-TX (PBS + 0.5% Triton-X) for 2×15min. Sections were then blocked for non-specific binding in a solution of 10% Normal Donkey Serum (017000121, Jackson Immuno) in PBS-TX for 1 hr at RT. After blocking, sections underwent an additional incubation in M.O.M. blocking buffer (1 drop M.O.M. IgG (MKB-2213-1, Vector Labs) per 4ml PBS) for 1 hr at RT, and then incubated in primary antibody overnight at 4°C in a solution of M.O.M. protein concentrate (BMK-2202, Vector Labs) in PBS. The next day, sections were washed 3×10min with PBS-T and incubated in relevant fluorescently-labeled secondary antibodies with M.O.M. protein concentrate (BMK-2202, Vector Labs) in PBS for 1 hr at RT. Sections were then washed 3×10min in PBS-T, mounted, dried, coverslipped with Gold Prolong Antifade Mounting Medium (Thermofisher P36930), and sealed with clear nail polish. For Thioflavin-S staining, mounted sections were stained with 0.015% thioflavin-S in 50% ethanol diluted in PBS for 10 min and washed three times for 5 min/wash with 1X PBS before coverslipping. Slides were imaged with an Aperio VERSA slide scanning microscope (Leica) at 10X magnification or a FV3000 confocal laser scanning microscope (Olympus) at 20X, 40X, or 60X.

Immunohistochemistry included the following antibodies: [Primary antibodies] Human Nuclear Antigen (ab215755, Abcam, 1:100); 3D6 (Elan Pharmaceuticals, 1:1000); AT8 (MN1020, Thermofisher, 1:100); MAP2 (PA1-10005, Thermofisher, 1:200); APOE (178479, Millipore, 1:1000); GFAP (Z0334, Dako, 1:500); GFAP (MAB3402, Sigma, 1:800); Iba1 (019-19741, Wako, 1:200); Iba1 (ab5076, Abcam, 1:100); Thioflavin-S (sc391005, Santa Cruz Biotech, 0.015% in 50% Ethanol/PBS); DAPI (62248, Thermofisher, 1:20000). [Secondary antibodies] Alexa Fluor; Jackson Immuno Research, 1:1000.

### Immunohistochemical Analyses

Immunohistochemical analyses were conducted in Fiji (ImageJ)^113^ via automated ImageJ macros to the extent possible. For all cell counts, images were set to a standardized threshold value across all conditions for each stain. For all aggregate counts, only particles within the hippocampus sized 10um^2 and above were analyzed. Aggregate densities were calculated and then normalized to transplant area to account for differing sizes of human cell transplants amongst conditions. For most quantifications, the results for each mouse represent the average of two hippocampal sections per mouse per stain. N for each condition is as follows: hE3-E3KI, 5 mice; hE3-E3KI-PLX, 6 mice; hE4-E4KI, 9 mice; hE4-E4KI-PLX, 9 mice; hEKO-E4KI, 4 mice; hEKO-E4KI-PLX, 6 mice. All compared confocal images were taken as z-stacks of similar depths and collapsed via z-project in Fiji (ImageJ). Analysts drawing regions of interest and setting standard threshold values were blinded to exclude possibility of bias. For all figures, unless otherwise specified, the following conditions were compared: hE3-E3KI vs hE3-E3KI-PLX, hE4-E4KI vs hE4-E4KI-PLX, hEKO-E4KI vs hEKO-E4KI-PLX, hE3-E3KI vs hE4-E4KI, hE4-E4KI vs hEKO-E4KI, hE3-E3KI-PLX vs hE4-E4KI-PLX, hE4-E4KI-PLX vs hEKO-E4KI-PLX.

### Microglia Isolation

Microglia were isolated using an adaptation of a previously described protocol^114^. The hippocampus of each mouse’s left hemi-brain was dissected on ice, rinsed with cold Dissection Buffer (1X HBSS, 5ug/mL Actinomycin, 10uM Triptolide, 27.1ug/mL Anisomycin), and placed into a pre-chilled 12-well plate with 2 mL/well of Hibernate A Buffer (BrainBits Hibernate A Buffer, 5ug/mL Actinomycin, 10uM Triptolide, 27.1ug/mL Anisomycin). Hippocampi from 4-6 mice per condition were pooled into one well for further processing. Tissue from each pooled sample was moved to a 100mm dish, minced with a razor blade (Personna double-blade, 0.004” thickness), then placed in a 5mL Eppendorf tube with 1mL pre-warmed Enzyme Buffer (1X DPBS with Ca^2+^and Mg^2+^, 1.5mg/ml Collagenase D, 50ug/ml DNaseI, 5ug/ml Actinomycin, 10uM Triptolide, 27.1ug/ml Anisomycin). Samples in Eppendorf tubes were then incubated in a 35°C water bath for 30 minutes, with gentle trituration at the 15-, 25-, and 30-minute timepoints. The homogenate was then filtered (70um MACS SmartStrainers) into a 50mL conical tube and centrifuged at 4°C at 450g for 7 minutes. The supernatant was carefully aspirated, and cells were resuspended in three parts 1X DPBS and one part Debris Removal Solution (Miltenyi Biotec). The homogenate was moved to a 15mL conical tube, and 4mL 1X DPBS was carefully overlaid atop the cell solution. After centrifuging at 3000g for 10 minutes at 4°C, the supernatant solution and myelin debris layer were carefully aspirated, and the cells were washed with PBS and spun down one last time at 1000g for 10 minutes before staining.

For staining, cells were resuspended in Blocking Buffer (1X DPBS, 0.2% BSA (Sigma), 2.5ug/mL CD16/CD32 (BD Biosciences)) and a portion of each sample was distributed into different tubes for staining controls. After a 5-minute incubation at 4°C, cells were incubated in Antibody Solution (1X DPBS, 0.2% BSA, 1.89ug/mL CD45 PE-Cyanine7 (Thermofisher), 3.53ug/mL CD11b Brilliant Violet 421 (BioLegend)) for 40 minutes at 4°C. Cells were then washed with 1X DPBS with 0.2% BSA, filtered through a 40um Flowmi Cell Strainer (Millipore Sigma) into a pre-chilled 5mL FACS tubes (Stem Cell Technologies), and kept on ice until sorting. DRAQ7 APC dye was added to each sample just before cell sorting to determine live/dead cells. Using a BD FACS Aria Fusion I or BD Aria II at 4°C with a 100um nozzle (20 psi), live single cells were identified and gated per standard sorting protocols, then further sorted to isolate a CD11b^+^/CD45^int^ microglia population (Supp Fig 6). Sorted cells were collected in pre-chilled 2mL LoBind tubes (Fisher Scientific) with Collection Buffer (RPMI 1640 HEPES Modified with 2% FBS (Fisher Scientific)). Cells were centrifuged at 500g at 4°C for 5 minutes, then loaded onto 10x Genomics Next GEM chip G at cell counts of approximately 3,000 cells (PLX-depleted conditions) to 25,000 cells (control and untransplanted conditions). The scRNA-seq libraries were prepared using the Chromium Next GEM Single Cell 3ʹ Library and Gel Bead kit v.3.1 (10x Genomics) according to the manufacturer’s instructions. Libraries were sequenced on an Illumina NovaSeq 6000 sequencer at the UCSF CAT Core.

To measure Csf1r levels, microglia from untransplanted E3KI and E4KI mouse hippocampi (6 mice per condition) were isolated and sorted as described above. Mice aged around 8 months were selected for this experiment to best measure Csf1r levels at an age comparable to that of chimeric mice during PLX3397 treatment. Hippocampi from each mouse were treated as separate samples, and all samples were sorted for CD11^+^/CD45^int^ microglia and measured for CSF1R levels using a BD LSR Fortessa™ X-20 cell analyzer.

Cell sorting and analyzing involved the following antibodies/dyes: Brilliant Violet 421^TM^ anti-mouse/human CD11b (Biolegend, 101236, 3.53ug/mL); CD45 monoclonal antibody (30-F11) PE-Cyanine7 (Thermofisher 25-0451-82, 1/89ug/mL); rat anti-mouse CD16/CD32 Mouse BD Fc Block (BD Biosciences 553141, 2.5ug/mL). DRAQ7 ^TM^ dye (Novus NBP2-81126).

### Pre-Processing and Clustering of Mouse scRNA-seq Samples

The scRNA-seq samples included a total of five samples, one from each of the different conditions (E3KI, E4KI, hE3-E3KI, hE4-E4KI, hEKO-E4KI). Each sample combined 4-6 hippocampi, one from each male mouse. The demultiplexed fastq files for these samples were aligned to the standard mouse reference genome (2020 version, <u>refdata-gex-mm10-2020-</u> <u>A</u>)^115^ separately using the 10x Genomics Cell Ranger v7.0.0 count pipeline, as described in the Cell Ranger documentation, and merged through CellRanger aggr. The include-introns flag for the count pipeline was set to true to count the reads mapping to intronic regions. For quality control assessment, cells filtered to keep only cells with greater than or equal to 250 UMI and 200 unique genes, and less than 5% mitochondrial genes. Normalization was performed using the NormalizeData and ScaleData functions of the R package for single-cell analysis Seurat v4.3.0.1^116–118^.

Graph-based clustering was performed using the Seurat v4.3.0.1 functions FindNeighbors and FindClusters. First, the cells were embedded in a k-nearest neighbor (KNN) graph (with k=20) based on the Euclidean distance in the PCA space. The edge weights between two cells were further modified using Jaccard similarity. Next, clustering was performed using the Louvain algorithm implementation in the FindClusters Seurat function. Clustering with 15 PCs and 0.5 resolution resulted in 16 distinct biologically relevant clusters, which was used for further analyses. Differentially Expressed Genes (DEG) were found using Seurat’s FindMarkers function. Volcano plots were created with EnhancedVolcano (Bioconductor package version 1.18.4) using unadjusted p-values, and the GO pathway analysis work was done with a combination of Enrichplot 1.18.4 and Clusterprofiler 4.6.2 (Bioconductor packages).

### Statistical Analyses

All immunohistochemical statistics were conducted as a two-way ANOVA in Graphpad Prism 10 (Graphpad Software Inc.). All data are shown as mean ± S.E.M. For all quantifications, unless specified, n represents the number of mice from which analysis images were gathered. No data were excluded based on statistical tests. p-values displayed are from the two-stage linear step-up procedure of Benjamini, Krieger and Yekutieli post hoc test for multiple comparisons (abbreviated in figures as Benjamini’s post hoc test for multiple comparisons). P <0.05 was considered significant, and all significant p-values were included in Figures or noted in Figure legends.

### Data Availability

The scRNA-seq datasets generated during the study will be made available at GEO (accession # xxxxxxxx) upon acceptance of the paper. Data associated with Figure 7 and Supplementary Figures 6 and 7 are also available in the Supplementary Information.

### Code Availability

All code generated during this study is accessible via reasonable request to the corresponding authors’ lab.

